# Single-Cell and Tissue-Specific CRISPR Editing Analyses Unveil New Insights to Off-Targets and Translocations

**DOI:** 10.64898/2026.04.13.718115

**Authors:** Alexandra Madsen, Niklas Selfjord, Marta Martinez-Lage Garcia, Anna-Lena Loyd, Gavin Kurgan, Mikaela Ståhlberg, Julia Lindgren, Julia Liz Touza, Leif Wigge, Mike Firth, Karl Nordström, Joseph Collin, Daniel Jachimowicz, Bastian Schiffthaler, Inken Dillmann, Panagiotis Antoniou, Aikaterini Emmanouilidi, Jonna Hellsten, Johan Forsström, Kerstin Magnell, Ashley Jacobi, Mark Behlke, Michelle Porritt, Katja Madeyski-Bengtson, Marcello Maresca, Pinar Akcakaya

## Abstract

CRISPR-Cas9 holds promise for treating genetic disease, but rare off-target mutations and structural variants remain as key safety concerns, especially at scales relevant to therapy. We established workflows to resolve Cas9 off-target activity *in vitro* at single-cell resolution and *in vivo* across different tissues. Using clonally expanded electroporated mouse embryos and embryonic stem cells, we reveal that individual cells exhibit unique off-target and translocation profiles, including events missed in bulk analyses. Integrating single-cell editing with chromatin accessibility, transcription, and DNA methylation measurements suggested that sequence-independent features modulate Cas9 access and cleavage, with preferential editing in regions characterized by open chromatin and lower methylation. In Cas9-inducible mouse models, editing analyses revealed organ-distinct off-target spectra, DNA repair pathway usage, indel patterns, and markedly varying translocation propensity between tissues. These findings demonstrate that off-target activity is heterogeneous across cells and context-dependent across organs, motivating sensitive single-cell analyses and organ-specific evaluation in preclinical development to more accurately assess risk and improve the safety of CRISPR-based genomic medicines.

## Introduction

CRISPR-based gene editing holds immense promise for treating human genetic disorders^1–4^, yet clinical translation requires rigorous identification and prevention of undesirable editing events, including off-target mutations and chromosomal aberrations^5–8^. The most widely used CRISPR nuclease, *Streptococcus pyogenes* Cas9 (SpCas9) scans double-stranded DNA for an NGG protospacer-adjacent motif (PAM) and uses a short guide RNA (gRNA) with an approximately 20-nucleotide spacer at the 5’-end to find its complementary target DNA^9^. Upon successful base-pairing between the spacer and the DNA target, Cas9 generates a double-strand break (DSB) that is repaired by endogenous DNA repair pathways. Imperfect DNA repair frequently yields small insertions and deletions (indels) at the target site^10, 11^. CRISPR nucleases can also introduce off-target DSBs at genomic loci with substantial sequence similarity to the intended target^6, 12–14^. Such mistargeting is a significant concern for therapeutic applications, as unintended mutations at off-target sites can be detrimental to normal cell functions and even potentially introduce oncogenic effects^15–18^. Moreover, multiple simultaneous DSBs in the genome can lead to chromosomal translocations or chromothripsis events, leading to compromised genomic integrity and unpredictable consequences^19, 20^.

Except for a few reports, principles that explain Cas9 genome-wide off-target activity other than sequence characteristics remain understudied. Genome editing outcomes are dependent on endogenous DNA repair pathways, and the utilization of the different pathways can vary greatly between different cell types and states^21–23^. A recent report showed that gene edited postmitotic neurons generate small non-homologous end joining (NHEJ)-related indels to a far higher degree than cycling iPSCs, which instead tend to generate larger, microhomology-mediated end joining (MMEJ)-associated deletions^22^. DSBs in neurons are resolved over a longer timeframe than cycling iPSCs, potentially due to slower rates of DNA damage response or a higher degree of indel-free repair.

Evidence on the link between chromatin state and off–target editing, especially *in vivo* where mismatches play critical role in impacting the expectedly low editing efficiencies, remains limited and in places conflicting. Most of the published CRISPR designs target coding regions with open chromatin. Early dead Cas9 (dCas9) binding studies showed mixed results, with some suggested epigenetic features such as CpG methylation and chromatin accessibility have little effect on targeting^24, 25^; whereas others found that chromatin accessibility is the strongest indicator of dCas9 binding^26, 27^. Studies with catalytically active Cas9 in doxycycline-induced reporter cancer cell lines have suggested that Cas9 binding and editing is significantly hindered by closed chromatin^28, 29^. One study using imprinted genes in mESCs found that heterochromatin can impede the kinetics of Cas9 cleavage but does not change the mutational outcomes^30^, while another report revealed that MMEJ is observed at higher frequency in specific heterochromatin contexts^31^. Tsai team found that in primary T-cells, bona fide off-target sites were 2-4 times more likely to be located in highly expressed genes, in open chromatin and at histone marks associated with active promoters and enhancers^32^. Similarly, the role of methylation on CRISPR binding and on-/off-target editing is unclear. Two early studies reported that DNA methylation status does not have a significant effect on Cas9 editing in human cells^24, 33^. A more recent report showed that high levels of promoter methylation, but not gene-body methylation, can cause a decrease in Cas9 activity and affect the ratio of mutational outcomes in plant cells, potentially through indirect changes in chromatin state caused by the methylation^34^. Collectively, these findings support a role for accessibility, but the magnitude and generality—especially for off targets in vivo—remain incompletely defined.

Considerable progress has been made in development of state-of-the-art tools to detect unintended editing events^32, 35–40^. For CRISPR-based therapeutics progressing towards the clinic, off-target safety assessments typically begin with a discovery phase in which multiple orthogonal methods are used to nominate candidate off-target sites. These methods include *in silico* algorithms^41–44^ that use sequence homology for off-target prediction, as well as experimental *in situ* or *in vitro* methods^32, 35, 36, 38, 39, 45–50^ that directly detect off-target DSBs in cells or in purified genomic DNA. Mutagenesis at off-target sites is subsequently evaluated in a validation phase using edited DNA from a therapeutically-relevant context. The most commonly used detection approaches are constrained by the current error rate of next-generation sequencing, which sets an arbitrary detection sensitivity of approximately 0.1%, or 1 in 1,000 cells^51^. In therapeutic genome editing, where billions of cells are typically targeted, this limitation risks missing low-frequency, yet clinically meaningful, CRISPR-induced mutations that may occur in millions of cells. Importantly, most off-target investigations to date have analyzed bulk cell populations or tissues^32, 38–40, 52–60^, potentially overlooking low-frequency mutations that fall below the detection threshold of conventional methods.

In our previous work, we have shown that careful design of gRNAs can minimize Cas9 off-target activity to levels undetectable by conventional NGS-based methods^52^, but also that off-target editing at levels below 0.1% can be detected at some of the same sites when using a more sensitive method such as Duplex Sequencing^40^. These results highlight the need for more sensitive and accessible off-target detection approaches in applications where very high safety standards are required.

We hypothesized that off-target editing is inherently heterogeneous and depends on unique properties of individual cells, such that even genetically identical cells that are edited at the same developmental stage may acquire distinct off-target profiles. Consequently, off-targets present in only a fraction of cells within bulk cell populations may evade detection because their frequencies fall below the sensitivity of conventional assays. By performing mutagenesis analyses at single cell level, we aimed to maximize the mutation detection in each analyzed individual cell and uncover rare off-target mutations that would otherwise be missed in bulk measurements. The analyses would most likely be unable to uncover all off-targets present in bulk populations but rather reveal some of the rare mutations present in bulk population, by maximizing their frequency in the particular analyzed individual cells in a randomized fashion.

In this report, we present a workflow for assaying CRISPR-Cas9 off-target mutations at the single-cell level in edited and clonally expanded single-cell mouse embryos and embryonic stem cells (mESCs). Our results demonstrate that individual cells exhibit distinct off-target profiles, even when edited simultaneously at the same cellular stage. Using a new computational algorithm, we were able to use the multiplex targeted sequencing data to quantify balanced translocations between on- and off-target sites, showing that individual cells display unique translocation products and that many of these products were undetectable when analyzing the bulk cell population. Furthermore, we have investigated the off-target propensity of different mouse cell types and tissues by using *in vitro* and *in vivo* inducible-Cas9 mouse models. As with edited single-cells, we could observe differences in off-target editing profiles and in the degree to which DSBs were resolved by micro-homology mediated end-joining in different cells and tissues. Collectively, these results highlight the complexity of CRISPR off-target activity, motivating the need for further research into factors beyond sequence homology that affect Cas9 editing at off-target sites and more sensitive off-target detection methods.

## Results

### Distinct off-target and translocation profiles in individual cells

To investigate our hypothesis, we established a single-cell editing analysis workflow (Fig. 1a) by delivering CRISPR RNP in a single cell suspension via electroporation followed by a 10–14-day single-cell expansion phase, providing sufficient DNA to capture the single-cell editing events using rhAmpSeq^61^ (Integrated DNA Technologies, IDT). RhAmpSeq is an amplicon sequencing method utilizing RNA-base-blocked primers and RNase H2 to enable high-fidelity multiplexing of hundreds of primer pairs into the same PCR reaction. This approach allowed us to analyze editing events at on- and off-target sites simultaneously in one-cell stage-edited mouse embryos, as well as mouse embryonic stem cells (mESCs). To only select the cells that were truly edited at single-cell stage for further analysis, we looked for one or two different on-target editing events at a frequency of 100% or 50%, indicating a clonal expansion from a single cell edited at either one or both alleles. We included two previously described single guide RNAs (sgRNAs) targeting the *Pcsk9* gene into our experiments: to be able to identify and study unwanted CRISPR off-target edits, we used the purposely promiscuous-designed “gP” control sgRNA that has many closely matched off-target sites (i.e. sites with a low number of mismatches to the sgRNA sequence) within the mouse genome and induced off-target editing at several of the analyzed sites *in vivo* in our previous study. In contrast, the highly specific “gMH” sgRNA that did not result in any detectable off-target editing in the same study was included to serve as a therapeutic example^52^. For each sgRNA, we designed a rhAmpSeq panel for off-target analysis consisting of off-target sites identified previously with the off-target nomination method CIRCLE-seq^37,52^. For gMH, the 69 off-target sites with the highest CIRCLE-seq score were included alongside 10 additional sites that were closely matched to the target sequence (i.e. carrying 3 mismatches relative to the target sequence) but were not identified by CIRCLE-seq. For the gP panel, the top 100 sites identified in CIRCLE-seq were included together with 22 additional sites assayed in the original publication^52^.

**Figure 1.**
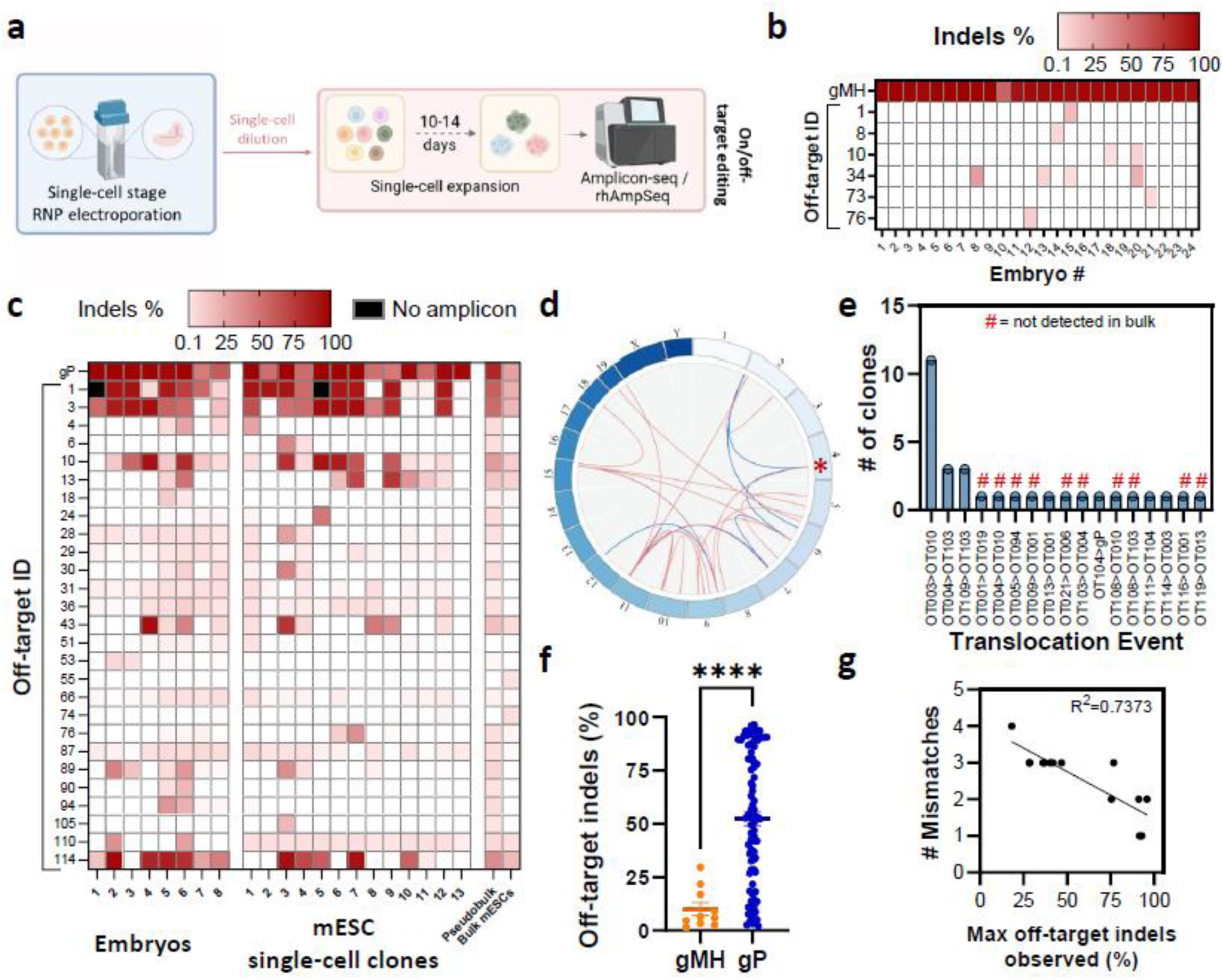
Single-cell editing reveals off-target editing differences between single cells. a. Overview of the experimental workflow for single-cell editing analyses in mouse embryos and mESCs. Created in BioRender. Madsen, A. (2026) https://BioRender.com/46i6ruz. b. Heatmap showing the percentage of indels detected at the indicated ”specific” gMH sgRNA on-/off-target sites in embryos. Embryos 1-10 have been expanded *in vivo*, embryos 11-24 *in vitro*. All off-target sites with indels detected at ≥0.1% in at least one clone are shown. c. Heatmap showing the percentage of indels detected at the indicated promiscuous gP sgRNA on-/off-target sites in embryos, single-cell edited mESC clones, and a pool of bulk-edited mESCs. Embryos 1-4 have been expanded *in vivo*, embryos 5-8 *in vitro*. Pseudobulk editing shows the mean editing calculated over all single-cell edited samples at each of the analyzed off-target sites. mESC bulk pool shows mean editing of three replicate pools 16 days after editing. All off-target sites with indels detected at ≥0.1% in at least one clone are shown. d. Circos plot showing gP sgRNA-induced translocations detected by PASTA in the embryos and in single-cell edited mESC clones. Blue lines indicate translocations that were also detected in edited cell pools, red lines indicate translocations that were only detected in the single-cell clones. The on-target site is marked with a red asterisk. e. Translocation analysis of rhAmpSeq with PASTA shows the number of gP-edited mESC clones in which the respective translocation was found. Translocation events marked with # were not detected in the bulk-edited mESC pools at either timepoint. f. Pooled visualization of the indels detected at gMH or gP off-target sites in clones derived from edited single cells shown in (b) and (c). Percentage indels in each clone are shown, as well as the mean±SEM. Mann-Whitney test, **** = p<0.0001. g. Correlation analysis between the observed indel percentage and the number of mismatches at gP off-target sites.

Using this workflow, we identified off-target mutations for the specific gMH sgRNA that we had initially reported to induce no Amplicon-seq-detectable off-targets in bulk tissues^52^, consistent with our previous report that identified off-targets when using the more sensitive Duplex Sequencing method^40^ (Fig. 1b, Supplementary Tables 1, 2). Moreover, each embryo and mESC clone edited with the promiscuous gP sgRNA exhibited a unique off-target profile (Fig. 1c, Supplementary Tables 1, 2), revealing divergent mutagenesis outcomes even when cells were edited simultaneously and at the same cellular stage.

The multiplexed nature of rhAmpSeq enables amplification of potential translocation products formed between the analyzed on-/off-target sites, permitting detection of such events using count-based statistical analyses, as had been previously shown^62^ (Fig. 1d, Extended Data Fig. 1a). We analyzed the translocation outcomes by leveraging the existing rhAmpSeq panel designed for the candidate off-target sites. Consistent with the off-target profiles, gP edited embryos and mESC single-cell clones displayed unique translocation profiles (Supplementary Table 3). None of the gMH-edited embryos or mESC clones showed translocations (data not shown). To note, this approach cannot detect unbalanced translocations because the resulting amplicons will generate non-sequenceable index configurations during library preparation (Extended Data Fig. 1b). Additionally, translocation products that fail to amplify due to technical factors, such as primer incompatibility or inefficient PCR, will not be detected even if present in the sample. Despite these limitations, the method provides important insights into translocation occurrence in a high-throughput manner when employing a promiscuous sgRNA and reveals unique translocation profiles in both embryos and mESC clones.

Next, we compared the gP off-targets and translocations found in single-cell mESC clones to the ones detected in bulk mESC culture that had been electroporated at the same time as the single cells and in a pseudobulk sample showing the mean editing from all individual cells at each of the analyzed off-target sites. We performed editing analysis in bulk mESCs at day 2 (timepoint of single-cell dilution) and at day 16 (timepoint of single-cell colony harvest). We uncovered several off-target mutations and translocation events in individual mESC clones that were not detectable in the bulk mESCs and pseudobulk sample (Fig. 1c - e, Extended Data Fig. 2, Supplementary Table 3). Off-target editing frequency and translocation events in the cell pools moreover declined significantly between day 2 and 16 after editing (Extended Data Fig. 2), indicating that edited cells might be lost during prolonged bulk culture due to potential impacts on fitness or proliferation.

We then compared the overall editing efficiency observed at off-target sites in gMH vs. gP-edited single-cells. Indel frequencies were significantly lower at gMH off-target sites than at gP off-target sites (Fig. 1f), suggesting that gMH off-target events occurred later during clonal expansion rather than at the one-cell stage. The higher off-target indel rate observed with gP may reflect differences in sgRNA-specific editing kinetics, implying that site-specific characteristics can slow Cas9 activity, consistent with previous *in vitro* reports^63, 64^. As expected, the off-target mutation frequency at gP sites negatively correlated with the number of mismatches within the spacer and PAM in single-cell clones (Fig. 1g). Because our data showed off-target editing occurs not only at the one-cell stage but also at later timepoints during the clonal expansion phase, to rule out the possibility of off-target mutations accumulating during clonal expansion rather than being generated by Cas9 nuclease, we established an adapted workflow employing whole-genome amplification (WGA) of single gP-edited mESCs to replace the clonal expansion step (Extended Data Fig. 3a). While this approach removes any potential bias introduced by the clonal expansion, it should be noted that the sequencing coverage of individual sites has a significantly higher inter-sample variation when using WGA DNA instead of genomic DNA of single-cell clones, greatly increasing the number of allelic dropouts during sequencing (Extended Data Fig. 3b). In addition, amplification bias and polymerase errors during amplification might further influence the editing outcome. Nonetheless, off-target editing analysis of the available data in WGA mESCs confirmed our previous findings, showing unique off-target profiles in different cells after editing with gP (Extended Data Fig. 3c). We additionally analyzed the presence of large structural variants in four of the WGA mESC samples in an exploratory manner by long-read sequencing, detecting inversions, large indels, duplications, and translocations in all samples (Extended Data Fig. 3d).

To test our workflow in a more therapeutically relevant setting, we repeated the single-cell expansion experiments in human hematopoietic stem and progenitor cells (HSPCs) using SpCas9 together with the previously described FANCF^35^, HEK4^35^, or CASGEVY^54^ sgRNAs or ePsCas9^65^ together with the HEK4 sgRNA. While no off-targets were detected in cell clones treated with the FANCF or CASGEVY RNPs (data not shown), the HEK4 sgRNA, designed to be promiscuous, introduced off-target edits at 16 of the analyzed sites, revealing unique off-target profiles also in the HSPC clones (Extended Data Fig. 4a, Supplementary Tables 1, 2). When using ePsCas9 nuclease, designed to induce less off-targets, together with the HEK4 sgRNA, we could detect off-target edits only at 6 of the analyzed off-target sites, confirming the higher fidelity of the enzyme (Extended Data Fig. 4a, Supplementary Tables 1, 2).

### Potential sequence-independent factors contributing to off-target editing

General principles that explain genome-wide off-target activity other than sequence characteristics remain largely unknown, with the exception of one report showing higher off-target activity on regions with open chromatin state and high transcriptional activity in bulk cell analysis^32^. To improve understanding of the mechanisms driving distinct off-target editing outcomes, we evaluated the target site accessibility at the time of CRISPR delivery in mock-electroporated mESCs by analyzing: 1) the cellular chromatin state using scATAC-seq and 2) gene expression using bulk- and scRNA-seq (Fig. 2a). Integrating the data with our gP single-cell off-target editing results, we observed an overrepresentation of coding regions compared to non-coding regions among the edited off-target sites (Fig. 2b). This observation is consistent with our previous investigation in bulk mouse tissues using the same promiscuous sgRNA^52^, confirming our earlier findings particularly for low-frequency off-targets detected at the single-cell level. A similar trend of preferential editing within coding vs non-coding regions was also observed for HSPCs edited with the HEK4 sgRNA (Extended Data Fig. 4b). Interestingly, exonic sites displayed a higher editing prevalence in regions characterized by open chromatin and lack of expression of the associated gene (Fig. 2c, d, Extended Data Fig. 5a, Supplementary Table 4), likely due to enhanced Cas9 complex accessibility to the target site. To ensure that the sites chosen for the off-target analysis are not inherently different from the other identified potential off-target sites, we additionally assessed the distribution of open chromatin and gene expression in all sites identified in CIRCLE-seq (Extended Data Fig. 5b), verifying absence of any significant outlier-effects. Moreover, our analysis of published mESC whole-genome bisulfite sequencing data sets^66, 67^ indicated a significantly lower average DNA methylation in a 1 kb-window around the cleavage location at edited sites compared to non-edited sites (Fig. 2e, Supplementary Table 5), suggesting an inhibitory effect of high DNA methylation on editing.

**Figure 2.**
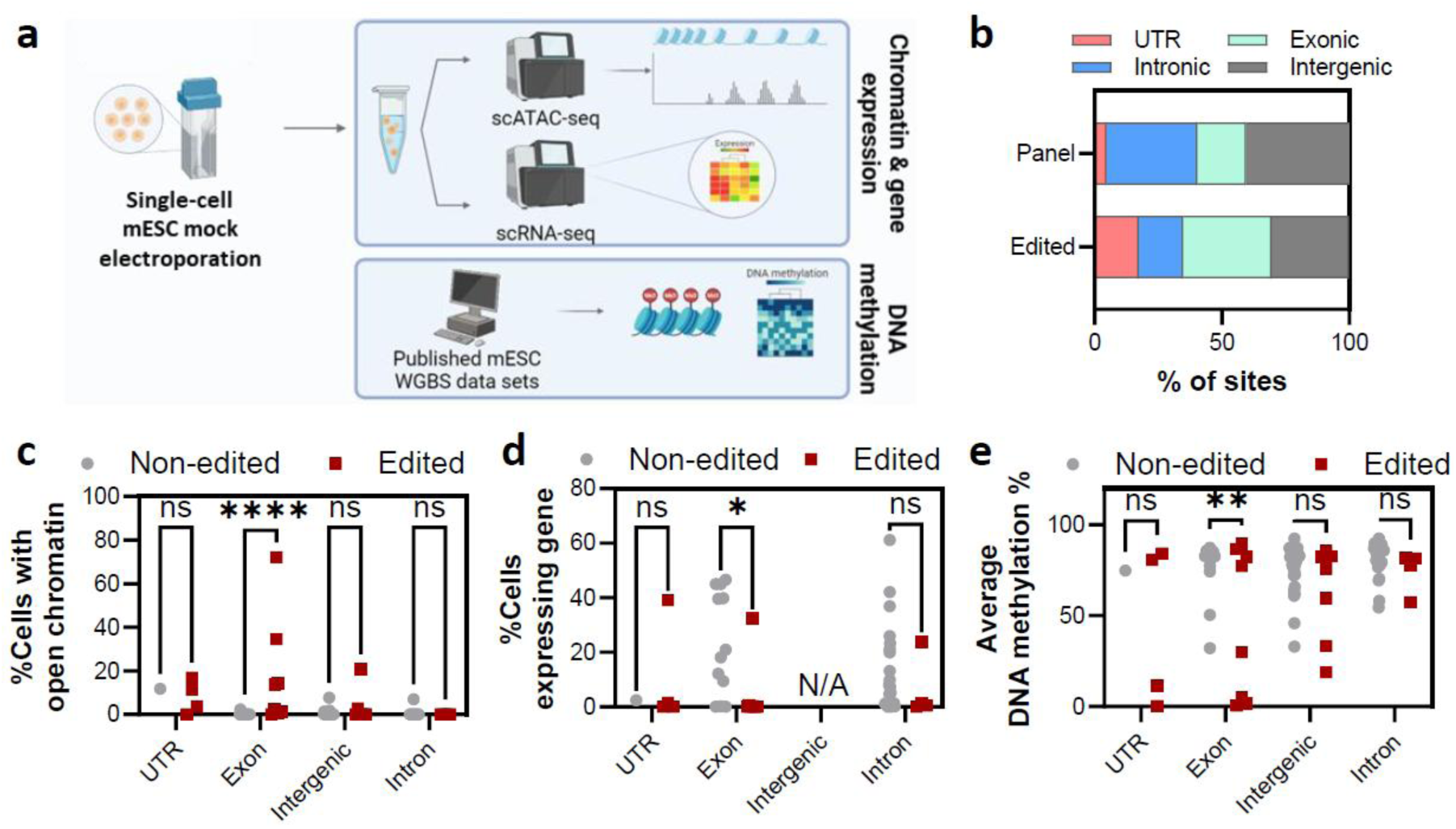
Differences in chromatin environment and gene expression at edited vs. non-edited sites. a. Overview of the workflow for analysis of chromatin environment and gene expression in mESCs. Created in BioRender. Madsen, A. (2026) https://BioRender.com/54olwqc. b. Bar plots showing the distribution of the analyzed gP on-/off-target sites over different functional genomic regions in all analyzed sites (“panel”) vs. in edited sites only (“edited”). c. Analysis of scATAC-seq data from mock-electroporated mESCs based on functional region of the respective gP off-target site. Plot showing percentage of cells with open chromatin at sites showing editing vs. sites that remain non-edited. Two-way ANOVA plus Sidak’s multiple comparisons test, **** = p<0.0001, ns = non-significant. d. Analysis of scRNA-seq data from mock-electroporated mESCs based on functional region of the respective gP off-target site. Plot showing percentage of cells expressing the off-target-associated gene at sites showing editing vs. sites that remain non-edited. Two-way ANOVA plus Sidak’s multiple comparisons test, * = p<0.05, ns = non-significant. e. DNA methylation analysis from whole-genome bisulfite sequencing data^66, 67^ at gP off-target sites showing editing vs. sites that remain non-edited. Plot showing the average methylation percentage of a region ±500 bp around each sgRNA off-target site. Two-way ANOVA plus Sidak’s multiple comparisons test, * = p<0.05, *** = p<0.001, **** = p<0.0001, ns = non-significant.

To test whether the aforementioned features could be used as predictor for off-target editing, we generated a new rhAmpSeq panel containing additional sites that were previously identified as potential off-targets in CIRCLE-seq discovery^52^. We classified these sites as likely to be edited (“editing panel”) or unlikely to be edited (“control panel”) based on our hypothesis (Supplementary Table 1) and analyzed these sites in the genomic DNA of the previously analyzed mESC clones. The editing panel consisted of 44 sites, however due to sequence complexities, only 12 sites reached the required coverage of >2000 sequencing reads, while ∼75% of sites remained below this coverage threshold and could not be analyzed. The control panel consisted of 47 sites, of which 46 sites had >2000 reads. Among the analyzed sites, we observed editing at a higher percentage of sites from the editing panel (30%) compared to the control panel (15%) (Extended Data Fig. 5c, Supplementary Table 2), suggesting the potential significance of including sequence-independent characteristics to the sgRNA design and prediction tools. We saw similar trends in chromatin, gene expression, and DNA methylation in edited vs. non-edited sites as in the initial gP panel (Extended Data Fig. 5d - 5f). It should, however, be noted that this analysis could be biased by the high number of sequencing dropouts in the editing panel due to sequence complexity at the analyzed sites. As none of the additional sgRNAs analyzed in our study induced a high enough number of off-target edits to enable a systematic analysis, we were not able to verify the broader translatability of our observations to other sgRNAs.

### Distinct off-target profiles in mESC-derived cell types and different organs *in vivo*

Considering the potential influence of chromatin environment, transcriptional activity and DNA methylation on off-target editing, we hypothesized that CRISPR might lead to diverse off-target editing outcomes in different cell types. To test this hypothesis, we generated a mESC line with doxycycline-inducible SpCas9 and constitutive promiscuous gP sgRNA expression, that enables editing after cell differentiation without having to rely on efficient delivery of the CRISPR reagents to the cells (Fig. 3a, Extended Data Fig. 6). The mESCs were then successfully differentiated into different cell types, representing each one of the three germ layers: astrocytes (ectodermal origin), cardiomyocytes (mesodermal origin), and hepatocytes (endodermal origin; Extended Data Fig. 7). Treatment with doxycycline, as expected, induced editing at the on-target site at a frequency between 45% and 80% in the different cell types (Fig. 3b). We analyzed off-target editing in the mESCs and the three mESC-derived cell types and observed that each of the cell types showed a distinct off-target profile (Fig 3b, Supplementary Table 6). In addition, analysis of shared edited sites across all cell types revealed cell type-specific differences in the extent of microhomology-mediated end-joining (c–MMEJ) usage during DNA repair and in the resulting indel profiles (Fig. 3c, d).

**Figure 3.**
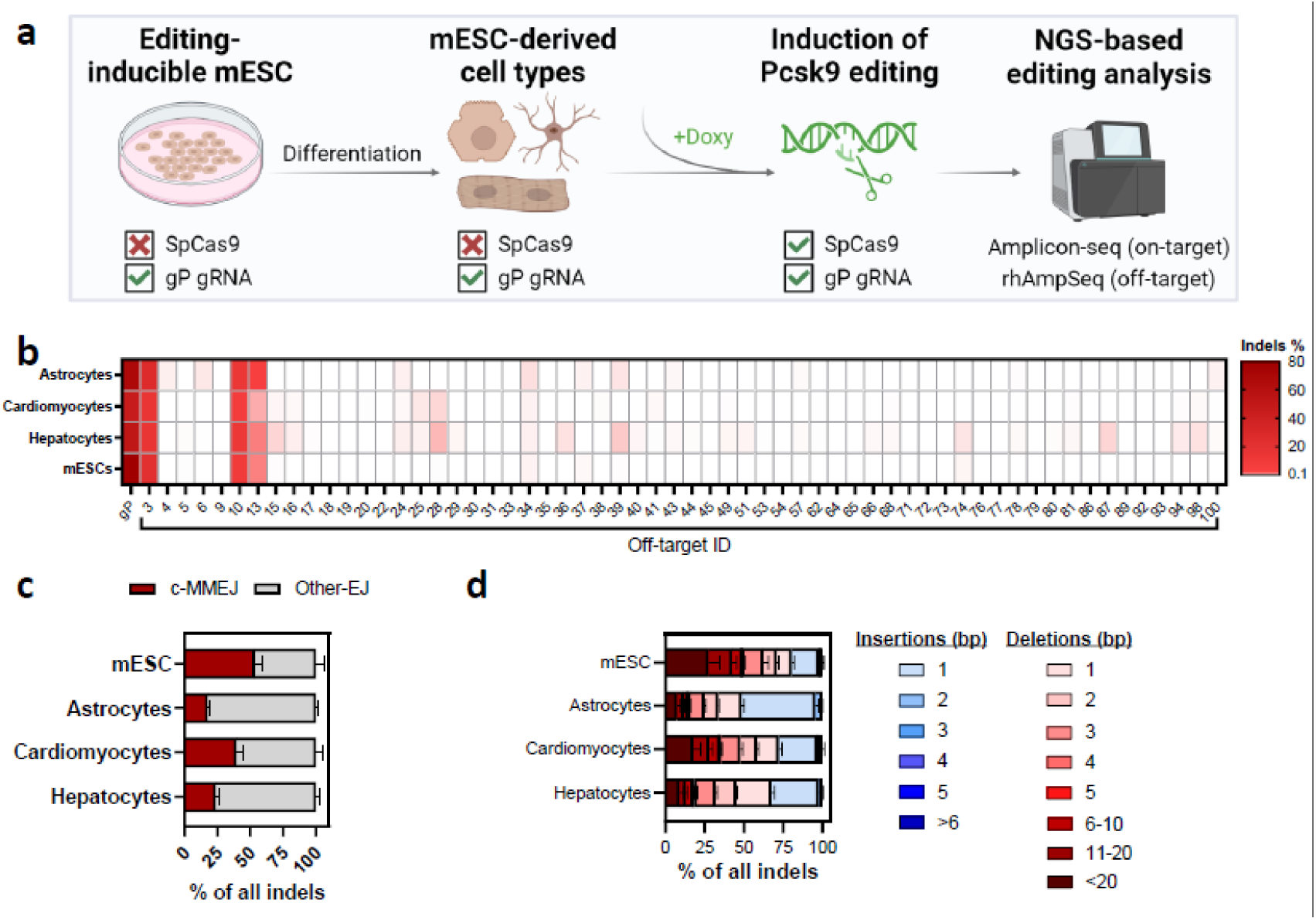
gP induces distinct editing outcomes in different cell types *in vitro*. a. Overview of the experimental set-up. Differentiation of the editing-inducible mESC to cardiomyocytes (mesodermal origin), hepatocytes (endodermal origin) and astrocytes (ectodermal origin), followed by SpCas9 induction with doxycycline and editing analysis. Created in BioRender. Madsen, A. (2026) https://BioRender.com/2oywezw. b. Percentage of indels detected at on-/off-target sites in different cell types after editing induction. Heat map showing the mean of n=4 replicates for each cell type. c. Comparison of DNA repair mechanisms at four sites that show indels in all cell types (i.e. on-target site, off-target sites 3, 10 & 13) upon doxycycline-induced SpCas9 expression with gP. n=4 replicates for each of the analyzed sites. c-MMEJ = micro-homology mediated end-joining; Other-EJ = other end-joining. d. Comparison of mutation profiles at four sites that show indels in all cell types (i.e. on-target site, off-target sites 3, 10 & 13) upon doxycycline-induced SpCas9 expression with gP. n=4 replicates for each of the analyzed sites.

To investigate the off-target profiles *in vivo* in different organs without the limitation of delivery success, we generated a mouse line (doxy-gP) derived from the doxycycline-inducible mESC line (Fig. 4a). We activated SpCas9 editing in mice via oral doxycycline administration, harvested different tissues after 12 days of doxycycline treatment and analyzed the editing outcomes at on-/off-target sites. While we could observe a low degree of leakiness of our construct (<10% on-target editing on average in non-induced animals), the resulting off-target editing efficiency in the absence of doxycycline was low enough to not interfere with the planned analysis (Supplementary Table 6). Consistent with our *in vitro* results, we observed distinct off-target profiles in various tissues upon SpCas9 activation (Fig. 4b, Supplementary Table 6), despite achieving similar SpCas9 expression and on-target editing levels (Extended Data Fig. 8a, b). Some of the off-targets were detectable in all organs, while others were detected only in a subset or single type of organ but with remarkably high reproducibility across the four biological replicates of each organ (Extended Data Fig. 8c). Similarly, at seven sites that were commonly mutated in all organs (i.e. on-target site, off-target sites 1, 3, 10, 13, 39 and 114) and in line with our *in vitro* observations, the different tissues exhibited clear differences in c-MMEJ utilization for DNA repair and indel profiles that were highly conserved within the four replicates of each organ, highlighting tissue-specific editing characteristics (Fig. 4c, d). For example, high frequencies of larger deletions (>6bp) were observed in lung, colon and spleen, whereas kidney, liver and pancreas showed preferential +1 insertions. Looking closer at the DNA repair underlying the detected indels at these seven sites, spleen, lung and colon showed >30% c-MMEJ DNA repair products at the investigated sites, while c-MMEJ accounted for only ∼7% of indel events in pancreas and kidney. Similar trends were observed upon analysis of all sites that showed >0.1% indels in the respective organs (Extended Data Fig. 9).

**Figure 4.**
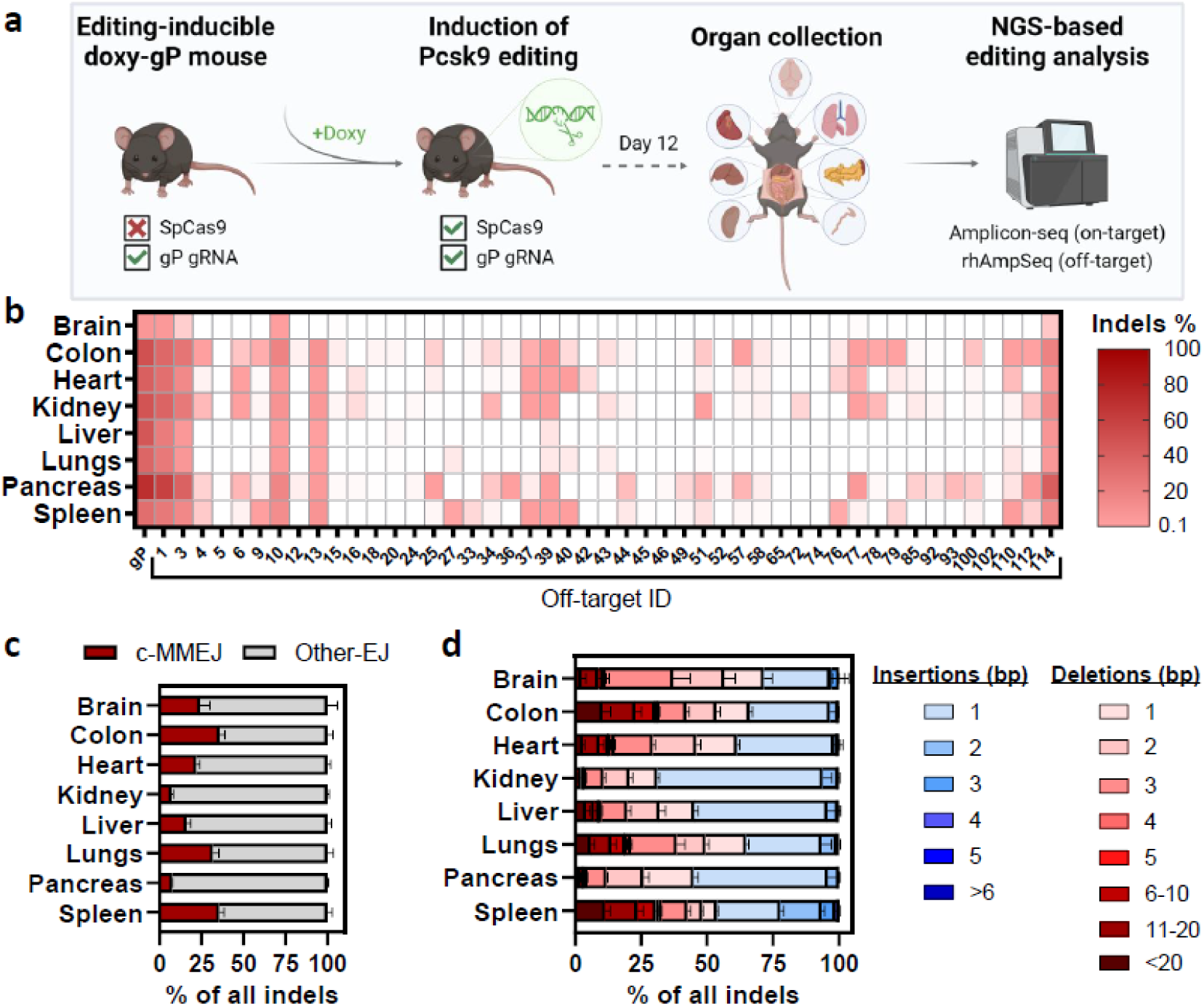
SpCas9 and gP promiscuous sgRNA induce distinct editing outcomes in different organs *in vivo*. a. Overview of the *in vivo* editing experiments using a doxycycline-inducible SpCas9-gP (doxy-gP) mouse model. Created in BioRender. Madsen, A. (2026) https://BioRender.com/xyz6iu5. b. Heat map showing the indel frequencies at the on-/off-target sites in different organs after editing induction by doxycycline treatment. Every data square presents the mean of n=4 organ replicates. All off-target sites with indels detected at ≥0.1% in at least one organ are shown. c. Comparison of DNA repair mechanisms at the seven on-/off-target sites commonly edited in the different organs upon doxycycline-induced SpCas9 expression with gP (i.e. on-target site, off-target sites 1, 3, 10, 13, 39 & 114). n=4 organ replicates for each of the seven analyzed sites. c-MMEJ = micro-homology mediated end-joining; Other-EJ = other end-joining. d. Comparison of indel profiles at the seven on-/off-target sites commonly edited in the different organs upon doxycycline induced SpCas9 expression with gP. n=4 organ replicates for each of the seven analyzed sites.

Building on our *in vitro* single-cell editing observations, we analyzed the presence of a potential relationship between exonic off-target editing and gene expression/DNA methylation *in vivo,* by performing bulk RNA-seq on tissues from the doxy-gP mouse line (Extended Data Fig. 10a, Supplementary Table 7) and analyzing published DNA methylation data sets^68–72^ in several organs of C57BL/6 and B6NCrl mice (Extended Data Fig. 10b). Unlike in the single-cell experiments, no clear correlation was observed between exonic off-target editing and expression/DNA methylation patterns in bulk cell populations *in vivo*.

### Organ-specific translocation propensity

We further investigated organ-specific DNA repair profile differences by interrogating translocation profiles across the different tissues. To normalize translocation frequencies to variable on-target activity, we derived a translocation score as the organ-specific cumulative translocation burden normalized by the on-target editing activity. The highest translocation scores were exhibited by heart, kidney, and pancreas, followed by lower translocation scores in colon, lungs, and liver, while brain and spleen displayed minimal to no translocation events (Fig. 5a, Supplementary Table 3). Each organ displayed unique translocations that were not found in the other organs (Fig. 5b). The presence of a specific translocation in different organs was however not predictable from the editing data, as exemplarily shown for two translocations that occurred only in the heart tissue in between two off-target sites, even though these two sites did not show the highest indel frequencies in heart (Fig. 5c), indicating additional factors influencing occurrence of translocations in a tissue-specific manner. Remarkably, the spleen stood out as the sole organ where neither shared translocations nor on-target/off-target translocations were found. These results suggest that translocation/off-target detection methodologies that rely solely on using on-target site as bait^36, 39^ may miss potential hits. As the occurrence of translocations depends on the NHEJ DNA repair pathway^73^, we analyzed the expression of genes involved in either NHEJ or MMEJ DNA repair in different organs but did not observe any correlation with the translocation score (Extended Data Fig. 11). Additionally, we performed *in vivo* editing experiments using doxycycline-inducible SpCas9 mice with constitutive expression of the specific gMH sgRNA (doxy-gMH) (Extended Data Fig. 12a). As expected, the analyses revealed no detectable off-target mutations or translocations in any of the analyzed organs (Extended Data Fig. 12b - e), verifying that the observed effects in doxy-gP mice were indeed editing-induced.

**Figure 5.**
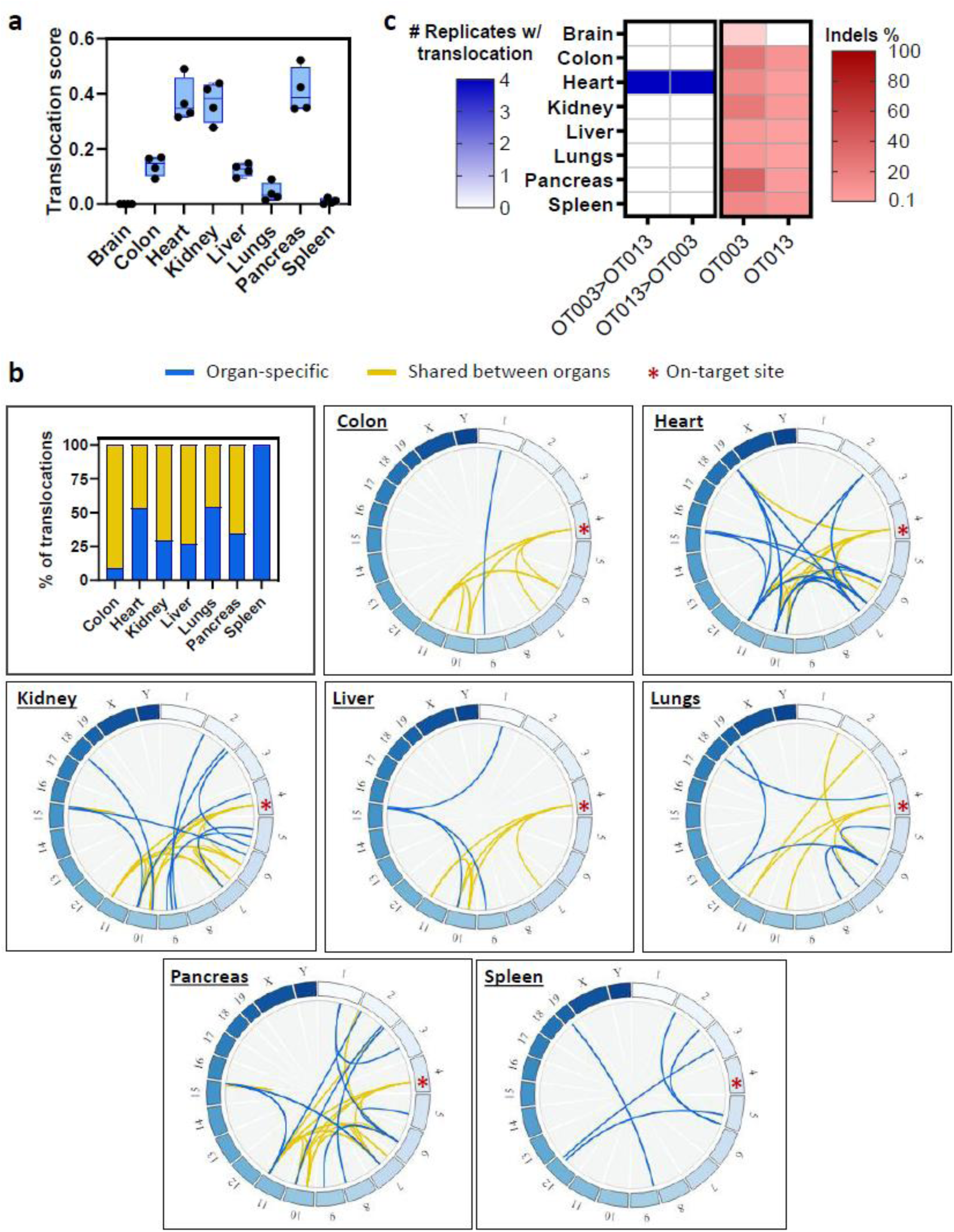
SpCas9 and gP promiscuous sgRNA induce distinct translocations in different organs *in vivo*. a. Comparison of the translocation score in different organs upon doxycycline-induced SpCas9 expression with gP. The translocation score was calculated by dividing the sum of all detected translocation frequencies to the on-target editing frequency for each organ. n=4 organ replicates per group. b. Overview of organ-specific translocations and translocations found in more than one organ. Circos plots depict the translocations detected for each organ, where blue lines indicate organ-specific translocations, yellow lines shared translocations, and the red asterisk marking the on-target location. c. Two representative heart-specific translocations in between two off-targets. Left heat map showing the number of replicates in which balanced translocation events between off-targets 3 and 13 could be detected in each organ. Right heat map showing the mean percentage of indels from n=4 replicates per organ at the two off-target sites that translocate.

## Discussion

Identification of unwanted editing events, such as off-targets and chromosomal aberrations, remains one of the biggest safety concerns for the development of CRISPR-based genomic therapies. We and others have made substantial progress in establishing tools to detect unwanted editing events^32, 38–40, 60, 74, 75^. However, most studies to date investigating off-targets have been performed in bulk cell populations^32, 38–40, 52–60^, potentially overlooking low-frequency mutations undetectable by conventional methods. The most commonly used off-target detection approaches are often limited by the current error rate and cost of next-generation sequencing, which set an arbitrary threshold for identifying mutations with frequencies lower than about 0.1%, 1 in 1000 cells. In the context of therapeutic genome editing in human liver that contains ∼200 billion hepatocytes^76, 77^, this means a risk of missing unwanted CRISPR mutagenesis that occurs in close to ∼200 million hepatocytes. We have recently developed a novel off-target detection approach that increased the detection sensitivity 10-fold using Duplex-seq^40^. Verifying the presence of mutations close to or lower than this detection sensitivity remains elusive. For well-designed therapeutic sgRNAs, this limitation may be insignificant, since very low-frequency mutations at safe genomic loci are unlikely to be harmful. However, when therapeutic efficacy depends on editing a specific sequence, such as correcting a disease-causing mutation, suitable sgRNA options may be limited. In these cases, comprehensive experimental evaluation of candidate sgRNAs is essential before proceeding, including not only thorough detection of unintended editing events but also rigorous assessment of their functional consequences that could lead to deleterious outcomes.

In this study, we established robust workflows and experimental models to address this gap, enabling us to perform comprehensive analyses of CRISPR editing outcomes in single cells and in different tissues *in vivo*, employing a specific (gMH) and a control promiscuous (gP) *Pcsk9*-targeting sgRNA^52^. Our main findings are: (1) Bringing the sensitivity to a single-cell resolution in an individual cell has revealed previously unidentified off-targets in bulk tissue^52^ and cell populations. (2) Even when cells are edited in parallel at the same cellular stage, each CRISPR-edited cell has a unique profile of off-target events and unwanted structural variations when treated with a promiscuous sgRNA. (3) While the sequence identity at the off-target site plays a large role in off-target recognition and editing, other sequence-independent factors such as DNA methylation, gene expression and chromatin status might influence the editing outcome at exonic sites. (4) The same sgRNA can cause entirely different off-target editing and translocations depending on which cell type it acts in.

We established a novel single-cell editing analysis workflow by taking advantage of the tightly controlled embryo electroporation system, by developing a robust single-cell stage edited clone selection approach based on the mutational profiles, and by using rhAmpSeq that enables simultaneous mutation analysis of hundreds of sites. We aimed to maximize detection within each profiled genome and uncover rare off–target mutations that are diluted in bulk. This approach would not reveal all off–targets present in the bulk population; rather, it reveals a subset of rare events by effectively “enriching” their allele frequency within the particular analyzed individual clones in a randomized fashion. Consistent with this rationale, our single–cell analyses identified off–targets that were missed in bulk cell populations despite deep coverage. Our results support the value of single–cell–based profiling for capturing rare or complex events that may evade bulk detection, while we continue to rely on well–powered bulk assays as a general conventional approach.

In line with what was described in a recent pre-print analyzing allelic editing outcomes at on-target sites in individual cells from a heterogeneous bulk cell population^23^ and in a recent study analyzing single-cell off-target events in ex vivo-engineered T cells^78^, the editing outcome in individual cells was unique and was not fully mirrored by off-target profiles generated from bulk cell analysis, with the latter missing several off-target editing events due to limitations in detection sensitivity with existing methods. This was especially apparent when analyzing cells edited with an sgRNA that was designed to be as specific as possible, similar to therapeutic sgRNAs, and which did not show any off-target editing *in vivo* in bulk analysis^52^. In line with our previous work revealing undetected edits when

increasing the detection sensitivity during off-target analysis^40^, editing analysis at single-cell level allowed detection of unwanted editing at six different off-target sites. However, the degree of off-target editing – both in terms of the number of sites showing editing and the kinetics of the edit compared to on-target editing – was significantly lower when using a specific-designed sgRNA compared to promiscuous sgRNAs, as expected. This underscores the importance and safety benefits of thorough sgRNA design for therapeutic purposes.

While we observed a moderate negative correlation between the number of mismatches between the sgRNA spacer and the off-target sites and the maximum observed off-target editing efficiency, not all editing events could be explained by sequence-dependent aspects alone. The accessibility of the genomic region harboring the off-target site appeared to be a major factor in governing off-target activity, as coding sequences with open chromatin being overrepresented in the sites showing off-target editing. This finding is in line with the observations of other research groups suggesting that closed chromatin negatively affects CRISPR activity in cells^29, 30, 79^. Additionally, we saw a significant negative impact of high gene expression on off-target activity in exonic regions, which might be caused by steric hindrance of the transcription machinery preventing Cas9 nuclease from binding the DNA. Another possibility is the fact that transcription-coupled DNA repair in general leads to a higher percentage of precise DNA repair of DSBs^80, 81^, which could mask Cas9 activity at these sites, as no detectable indels are created in the process. Similarly, the negative impact of DNA methylation on editing might be due to steric hindrance/binding competition at the respective genomic sites, especially impacting the lowly edited sites as previously shown^82^ – which makes it even more important for lowly edited off-target sites. Our validation panel supports the concept that incorporating sequence–independent features can improve off–target prediction beyond sequence similarity alone. Sites prospectively classified as “likely edited” showed roughly double the hit rate versus controls (30% vs 15%) and recapitulated chromatin, expression, and methylation trends observed in our initial panel, suggesting these contextual features are generalizable. However, the substantial sequencing dropouts in the editing panel due to sequence complexity limit definitive conclusions, underscoring the need for higher–coverage assays or orthogonal methods to robustly quantify predictive gain and to fully elucidate the underlying mechanism. As we identified very few exonic off-targets for the additional sgRNAs analyzed in this study, we were not able to perform a powerful analysis to verify the universal generalizability of our findings. Nonetheless, our observation that exonic regions with low DNA methylation and low gene expression were preferentially edited in our experiments adds important new insights to the growing body of evidence for the involvement of these factors in regulating Cas9 activity^34, 82^.

Building on the hypothesis of genomic context influencing the editing outcome, we compared the editing outcome in different cell types and organs *in vivo*. Since there are substantial differences in the transfectability of different cell types both *in vitro* and *in vivo* that could bias the editing analysis^83^, we generated an inducible model controlling the expression of SpCas9 with a doxycycline-inducible promoter paired with constitutive expression of the sgRNA in all organs. This model allowed us to reach similar expression of the editor in all analyzed organs *in vivo* with the exception of the brain, where the blood-brain barrier prevented efficient distribution of doxycycline to the tissue. Using this model, we confirmed our hypothesis that the editing outcome using the same sgRNA differs greatly depending on which organ/cell type is edited, observing organ-specific off-target profiles, c-MMEJ DNA repair utilization, indel patterns, and translocation propensity. It is worth keeping in mind however, that a specific translocation might – depending on the cell type in which it arises – affect cellular fitness of the cell, leading to cell death and subsequent non-detectability of the translocation in an organ-specific manner^84^. While some of the detected translocations might indeed be the product of organ-specific editing and translocations events, others might consequently simply be lost over time in a subset of organs, giving the impression of organ-specific translocation occurrence. Further research employing time course experiments will be needed to fully understand these intriguing results. Together, our *in vivo* findings have important implications for the safety of therapeutic gene editing medicines, as they highlight the importance of performing editing studies in the target cell type rather than using surrogate cell lines for more predictive outcomes reflective of the therapeutic approach, as well as the significance of stringent biodistribution analyses coupled with organ-specific editing analysis in both on- and off-target organs.

Following our observations on potential involvement of genomic context on editing outcome, we analyzed the observed off-target activity in the context of gene expression activity and DNA methylation in the different organs. However, the results did not replicate our *in vitro* findings, showing no clear correlation of off-target editing with gene expression or DNA methylation status *in vivo*. One potential reason for these differences is the use of highly-proliferative, undifferentiated cells in the *in vitro* setting, which might display drastically different attributes in the analyzed genomic factors compared to the fully differentiated adult cells in our *in vivo* experiments^85, 86^. Additionally, the *in vivo* analysis has the major limitation of both the off-target and genomic context analysis being done in bulk tissues. As expected, and as recently shown in the context of on-target editing^23^, bulk analysis results are biased towards the most abundant cell type within the tissue. Since it was not possible to control for the exact cell composition in the analyzed tissue samples in both our experiments and the published data sets, making direct correlations between the observed editing and the genomic context remains difficult. The presence of multiple different cell types with diverging chromatin environment and gene expression at the analyzed regions might therefore mask any potentially present effects. Future studies will need to employ extensive parallel analysis of editing outcomes and genomic context on a single-cell level across a large number of sgRNAs to fully understand the interplay between all these different factors and allow better prediction of unwanted editing outcomes in different target organs and cell types.

In summary, our findings represent one of the first robust demonstrations that CRISPR-Cas nucleases induce distinct off-target mutation profiles and translocations in single cells *in vitro* and in various tissues *in vivo*. Our analyses confirm previous discoveries regarding the influence of genomic location and chromatin accessibility on off-target editing at the single-cell level. Additionally, we provide new mechanistic insights, highlighting factors such as DNA methylation and gene expression at exonic sites, which may affect accessibility to CRISPR and off-target editing. We recommend that careful sgRNA design should consider not only sequence context but also factors influencing the genomic accessibility to help minimize off-target mutations. Our findings highlight the importance of performing the editing studies in the target cell type rather than using surrogate cell lines for the most predictive outcomes reflective of the therapeutic approach. We believe that our single-cell editing analysis methodology introduces a complementary strategy to the existing off-target detection methodologies for candidate therapeutic sgRNAs that may pose a higher risk of off-target activity and therefore warrant more extensive profiling, especially as it is applicable to clinically relevant mitotically active cells like T cells or hematopoietic stem cells, regardless of the nuclease choice. We anticipate that the collective results and methods described here will strongly motivate the development of more sensitive off-target detection methods to identify even the rarest CRISPR editing events, facilitating thorough risk assessment of gene editing outcomes, and ultimately fostering the development of safer genome editing therapies.

## Online Content

### Methods

#### Mouse embryo electroporation and expansion

Cryopreserved C57BL6/N mouse zygotes were purchased from Janvier labs (#SE-ZYG-CNG) and used for ribonucleoprotein (RNP) complex electroporation. The RNP complex was formed by mixing 6 µM SpCas9 protein (Alt-R SpCas9 Nuclease V3, #1081059, IDT) with 6.6 µM sgRNA (IDT) in Cas9 buffer (0.5 M KCl, Sigma 60142 0.1 M Hepes, 1 M ph 7.2-7.4; Life Technologies #15630) and kept at room temperature for 10 minutes. Two different guide RNAs were used, either the promiscuous gP or the specific gMH sgRNA. 50 zygotes/shot were placed in a 1 mm BIORAD GenePulser®Micropulser cuvette (BioRAD #165-2089), containing 10 µl Opti-MEM (Gibco™Opti-MEM™; Thermo Fischer Scientific #31985062) and 10 µl RNP complex. The BIORAD Gene Pulser Xcell Electroporation system was used to electroporate zona intact embryos using the following settings: square wave protocol, voltage 30V, pulse length 3 ms, number of pulses 10, pulse interval 100 ms, cuvette length 1 mm. After electroporation, 100 µl KSOM+AA was used to transfer zygotes from the electroporation cuvette to prewarmed KSOM in embryo dishes. The zygotes were left to recover in a water jacket incubator at 37 °C and 5% CO_2_ over night. Zygotes that developed into two cell stage embryos were either implanted into pseudopregnant B6D2F1/Crl foster mothers (gMH embryo 1-10, gP embryo 1-4) or transferred to agarose-coated 24 well cell culture dishes and grown in KSOM for *in vitro* blastocyst formation (gMH embryo 11-24, gP embryo 5-8). Mouse embryos from *in vitro* and *in vivo* expansions were collected after 10-14 days.

#### mESC culture and RNP editing

Primogenix C57Bl6/N (PrX) mouse embryonic stem cells (mESC) were cultured on plates coated with 0.1% gelatin (Stemcell Technologies) in basic mESC growth medium (Knockout DMEM (Gibco) supplemented with 15% FBS (Gibco), 1% Pen/Strep (Gibco), 0.1 mM beta-mercaptoethanol (Gibco), 2 mM Glutamax (Gibco), 1x MEM NEAA (Gibco), 1000 U/ml LIF (Sigma-Aldrich)). For electroporation, cells were harvested by trypsinization and electroporated with an RNP complex of 100 pmol SpCas9 protein (IDT) and 100 pmol synthetic sgRNA (IDT) using the P3 4D-Nucleofector X kit (Lonza) and the Amaxa electroporation system (Lonza) with program CG-104. Cells were allowed to recover for 48h on gelatin-coated cells, before being subjected to single cell-seeding for clonal expansion or sorted as single cells into a 96 well plate using the BD FACSAria III cell sorter (BD Biosciences). Additionally, the remaining electroporated cell pool was cultured for DNA isolation on day 2 and day 16 after electroporation to serve as a bulk cell population for comparison to single-cell editing data.

#### HSPC culture, ribonucleoprotein transfection and colony-forming cell (CFC) assay

Granulocyte colony-stimulating factor (G-CSF)-mobilized peripheral blood CD34^+^ HSPCs from healthy donors were purchased (Charles River). Written informed consent was collected from all adult subjects. HSPCs were thawed and cultured at a density of 500,000 cells/mL in StemSpan medium (STEMCELL Technologies) enriched with 100 U/ml penicillin/streptomycin (Gibco), 2 mM L-glutamine (Gibco), 250 nM StemRegenin1 (STEMCELL Technologies), 20 nM UM729 (STEMCELL Technologies), and supplemented with the following recombinant human cytokines (PeproTech): human stem cell factor (SCF; 300 ng/mL), Flt-3L (300 ng/mL), thrombopoietin (TPO; 100 ng/mL), and interleukin-3 (IL-3; 60 ng/mL). Cells were maintained in culture for 2 days before proceeding with nucleofection.

Ribonucleoprotein (RNP) complexes were assembled at room temperature using either 4.5 µM SpCas9 (IDT) or 4.5 µM ePsCas9^65^ in combination with 20 µM synthetic sgRNA (IDT). HSPCs, at a density of 200,000 cells per condition, were transfected with these RNP complexes using the P3 Primary Cell 4D-Nucleofector X Kit (Lonza) and the CA-137 program on the Nucleofector 4D device. Mock transfected cells served as negative controls for the experiments.

Post-nucleofection, the HSPCs were plated at a concentration of 500 cells/mL in a methylcellulose-based medium (GFH4435, STEMCELL Technologies) conducive to erythroid and granulo-monocytic lineage differentiation. BFU-E and CFU-GM colonies were analyzed 14 days after plating. Individual BFU-Es and CFU-GMs colonies were randomly selected and collected to assess on-targeting efficiency through NGS-based amplicon sequencing and examine off-target editing using rhAmpSeq panels.

#### DNA isolation and next generation sequencing (NGS) analysis

DNA for editing analysis was extracted with one of the following methods: (a) for *in vivo*-expanded embryos, the Gentra Puregene kit (QIAGEN) was used; (b) *in vitro*-expanded embryos were processed with the PicoPure DNA extraction kit (ThermoFisher); (c) DNA from mESC clones and pools was extracted with QuickExtract DNA extraction solution (Lucigen). All procedures followed the respective manufacturer’s instructions.

On-target editing for gP and gMH guides was analyzed by NGS-based amplicon sequencing as described previously^40^. In short, amplicons of the on-target loci were generated with Phusion Flash High-Fidelity Mastermix (ThermoFisher) using target specific primers (see Supplementary Table 8) including adapters for subsequent index incorporation. After PCR purification using Ampure XP beads (Beckman Coulter), unique Illumina indexes were introduced to the amplicons using KAPA HiFi HotStart Ready Mix (Roche). The PCR products underwent a second round of bead purification prior to sequencing on an Illumina NextSeq system according to manufacturer’s protocol. NGS data was demultiplexed with bcl2fastq software. The resulting fastq files were aligned to the expected target reference sequences using bwa mem (0.7.15-r1140)^87^. Variants were identified from the SAM files generated using a custom python script based on the pysam package. Editing outcome and mutation pattern analysis was done using RIMA as described previously^88^. Successful single-cell editing in embryos and mESC clones was confirmed by analyzing both the on-target editing efficiency (∼50% or ∼100% for mono-allelic and bi-allelic editing, respectively) and the on-target mutation pattern (presence of 1 or 2 distinct mutations for homozygous and compound-heterozygous edits, respectively).

Off-target editing was analyzed by multiplexed rhAmpSeq^61^ using the rhAmpSeq Library Kit (IDT) with a set of customized rhAmpSeq panels (see Supplementary Tables 1, 2 and 6) according to standard protocol. The panel for gMH and gP panel 1 included the sites originally analyzed in Akcakaya & Bobbin et al.^52^. In addition, a second panel comprising the 100 sites with highest gP *in vitro* cleavage scores was designed (gP panel 2). rhAmpSeq libraries were sequenced on the Illumina NextSeq system according to manufacturer’s protocol and analyzed as described above. Mutation frequencies of untreated and treated transgenic mice were normalized to their respective wild-type counterparts to remove any sequencing-related errors. Sites reaching less than 2,000x coverage were excluded from the analysis and marked in the heat maps as drop-outs (“no amplicon”). Only mutations >0.1% after this background subtraction were considered relevant. Statistical analysis of off-target editing for non-single-cell experiments was performed as described previously^89^. P values were calculated with MASS package in R statistics by fitting a negative binomial regression and with the logarithm of the total number of reads as the offset to the control and edit samples for each evaluated site. The results were adjusted for multiple comparisons using the Benjamini and Hochberg method (function p.adjust in R version 3.6.0). Indels in doxycycline-treated replicates were considered significant if mean indel frequency >0.1, p < 0.05, and nuclease-treatment coefficient > 0.

#### Translocation analysis with PASTA

To quantify translocations from rhAmpSeq data, Primer Anchored Statistical Translocation Analysis (PASTA) was used (manuscript in preparation). Briefly, expected primers were identified in reads using fg-idprimer (https://github.com/fulcrumgenomics/fg-idprimer<u>; -k=6, -K=8, -S=5, --max-mismatch-rate=0.07</u>). Following this, treatment/non-treatment pairs had their counts paired and primer count frequencies subjected to a one-tailed hypergeometric test with Benjamini-Hochberg correction (statsmodel v0.15.0; default settings) to calculate an adjusted p-value (p^adj^). Unexpected primer pairs with p^adj^ < 0.01 with no flags were classified as a translocation and had the translocation frequency (P) calculated using the following equation:

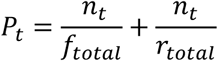

Where ‘n’ is equal to the count of the unexpected primer pair of interest, ‘t’ is the significant translocation being interpreted, ‘f’ is the total count of the shared forward primer events excluding the count participating in the ‘n’ translocation event, and ‘r’ is the total count of shared reverse primer events excluding the count participating in the ‘n’ translocation event. The translocation frequency is then adjusted by the background level frequency in the control by subtracting any translocation frequency observed in the control sample from the treatment frequency. Total translocation burden (B) was calculated using the following equation:

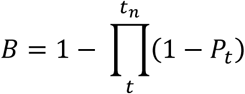

Where ‘t’ is equal to a significant translocation, and ‘t_n_’ is equal to the last significant translocation of all translocations. All translocations for the purposes of this equation are assumed to be occurring independently. Translocation analyses were performed for the gMH panel and gP panel 1 only.

#### Droplet digital PCR (ddPCR)

For the targeted analysis of translocations in mouse embryos, custom FAM-labelled ddPCR-assays were designed to target the translocation event where the 3’ end of OT001 is fused to the 5’ end of OT114 (balanced translocation) as well as where the 5’ end of OT001 is fused to the 5’ end of OT114 (unbalanced translocation). HEX-labelled Ap3b1 assay (Bio-Rad, dMmuCNS801070401) was used as reference. Sequences for the primers and probes are listed in Supplementary Table 8. ddPCR reactions were prepared to each contain 1x ddPCR Supermix for Probes (no UTP) (Bio-Rad), 1x FAM-labelled custom translocation assay, 1x HEX-labelled reference assay, 5 U HindIII (NEB), and 50 ng genomic DNA. An automated Droplet Generator (BioRAD) was used to generate droplets prior to PCR amplification in a C1000 Touch™ Thermal Cycler (Bio-Rad) using the following cycling conditions: 95 °C 10 min, 40 cycles of [94 °C 30 s, 59 °C 1 min and 72 °C 1 min] and 98 °C 10 min. The droplets were read on the QX 200 Droplet reader (Bio-Rad) using ddPCR™ Droplet Reader Oil (Bio-Rad). QuantaSoft software (BioRad) was used for analysis with manual fluorescence amplitude threshold setting based on average amplitude of FAM and HEX channels in edited samples and negative controls. The same threshold was applied to all wells of the ddPCR-plate using the same translocation assay.

#### Whole genome amplification for editing analysis

Whole genome amplification (WGA) of single-cell sorted mESCs was performed using the REPLI-g Single Cell Kit (QIAGEN). The WGA DNA was then purified using ethanol precipitation. WGA success was confirmed using the Fragment Analyzer DNF–467 Genomic DNA 50 kb Kit (Agilent Technologies). In addition, DNA from the remaining electroporated cell pool was extracted using QuickExtract DNA extraction solution (Lucigen). All procedures were performed according to the manufacturer’s instructions.

#### Long-read nanopore sequencing for structural variant analysis

WGA products from single mESCs were treated with T7 Endonuclease I (NEB) to remove branched structures, then subjected to size selection for long fragments using AMPure XP beads (Beckman Coulter). Both procedures were performed according to the Ligation Sequencing gDNA V14 — Whole Genome Amplification protocol (SQK-LSK114, Oxford Nanopore Technologies). Minor adjustments were introduced for bead–based purification: the custom buffer was prepared from PEG 8000 (NEB), 0.5 M EDTA (pH 8.0; Thermo Fisher), 5 M NaCl (Sigma–Aldrich), 1 M Tris (pH 8.0; Thermo Fisher), and nuclease–free water (Thermo Fisher). Bead-purification was conducted using either the custom buffer specified in the protocol, or the standard buffer supplied with AMPure XP beads. The concentration of whole genome amplified DNA was measured using Qubit High Sensitivity Quantification Kit (Invitrogen, Thermo Fisher Scientific). DNA fragment size distribution was evaluated on a 12-capillary 5200 Fragment Analyzer using the Genomic DNA 50 kb assay (Agilent Technologies). For each sample, 1000 ng of input DNA was used to prepare libraries with the Ligation Sequencing Kit V14 (SQK-LSK114, Oxford Nanopore Technologies) following manufacturer’s instructions. 250 ng of the final library per sample (estimated ∼50 fmol of DNA molecules) was loaded onto separate R10.4.1 flow cells (FLO–PRO114M, Oxford Nanopore Technologies) and run on a PromethION 2 Solo for 92 hours.

Basecalling was performed using dorado in SUP mode to ensure maximum per-base sequence quality. The GRCm38 (mm10) primary assembly was used as the reference genome throughout all analyses^90^. Gene-level annotations were obtained from GENCODE release vM25^91^. Raw reads were aligned to the GRCm38 reference genome using minimap2 v2.29^92^ in long-read high-quality mode (-ax lr:hq). Resulting alignments were coordinate-sorted and output in CRAM format using samtools sort (SAMtools v1.22)^93^, and CRAM index files were subsequently generated with samtools index. Per-sample alignment statistics were computed with samtools stats v1.22^93^. Summary fields were extracted from the SN section of each report to obtain the total number of mapped reads and the total number of CIGAR-corrected mapped bases. Genome-wide base coverage was estimated by dividing the number of mapped bases by the size of the GRCm38 genome assembly (2,730,871,774 bp), and the fraction of mapped reads was calculated as the ratio of mapped reads to the sum of all mapped and unmapped reads. Both metrics were inspected across all samples to confirm adequate sequencing depth and alignment efficiency prior to variant calling. Structural variants (SVs) were called independently for each sample using Sniffles2 v2.6.2^94^. Each CRAM file was processed with tandem-repeat annotations for mm10 (TRF BED format) supplied via the --tandem-repeats argument to improve breakpoint resolution in repetitive genomic regions. A per-sample Sniffles Fingerprint (SNF) file was generated during this step to enable subsequent joint genotyping. Per-sample VCF files were additionally produced by re-running Sniffles2 in single-sample mode using the respective SNF files as input. Output was written to BCF format and indexed using bcftools v1.22^93^. Joint SV genotyping across all four samples was performed with Sniffles2 v2.6.2^94^ in population mode, with all four SNF files provided simultaneously as input. Tandem-repeat annotations and the GRCm38 reference sequence were supplied consistently across all Sniffles2 invocations. The resulting multi-sample VCF file was used as the basis for all downstream filtering and annotation steps. The multi-sample VCF was imported into R using the VariantAnnotation Bioconductor package^95^. Two quality-based criteria were applied conjunctively to retain high-confidence SVs. First, a genotype quality (GQ) filter required that at least three of the four samples carried a GQ score exceeding 30. Second, a genotype (GT) filter required that at least three of the four samples were assigned a well-defined diploid genotype, defined as homozygous reference (0/0), heterozygous (0/1), or homozygous alternate (1/1). Only variants satisfying both conditions simultaneously were retained for downstream analysis. Filtered SVs were annotated with overlapping gene models from the GENCODE vM25 GFF3 annotation file^91^, which was imported using the rtracklayer package^96^. Genomic overlap between SV breakpoint intervals and gene features was computed with GenomicRanges^96^. For each SV type (deletions, insertions, duplications, inversions, and breakend events), the resulting annotated tables were exported to Microsoft Excel format using the openxlsx R package. Read alignments within a 5,000-bp window centered on the on-target CRISPR edit site (chr4:106,463,846–106,463,865) were extracted from each CRAM file using samtools view and visualized as alignment tracks using the Gviz Bioconductor package, with the on-target interval highlighted.

#### scATAC-seq/RNA-seq

The scATAC/RNAseq experiment was performed on mock-electroporated mESCs using the Chromium Next GEM Single Cell Multiome ATAC + Gene Expression kit (10x Genomics), according to the manufacturer’s recommendations, and targeting a recovery of ∼5000 cells. The libraries were sequenced on a NextSeq 2000 (Illumina). RNA-seq and ATAC-seq data were demultiplexed individually using bcl2fastq (v2.20.0.422). The resulting fastq files were processed using 10x Genomics Cellranger (cellranger-arc count v 2.0.2). The peak and gene count matrices and fragments file from Cellranger were used as input for analysis in R (4.2.3) with Signac (1.10.0)^97^ and Seurat (4.3.0.1)^98^. The dataset was filtered into high quality cells, keeping cells with >1000 total fragments, percent reads in peaks >50 and corresponding gene expression count >1000. After filtering, peaks were re-called using MACS2 (2.2.9.1)^99^ wrapped in the CallPeaks function of Signac and a final peak count matrix was generated. For each off-target site with an overlapping peak, the fraction of cells where this peak was detected (peak count >0) was calculated. For each off-target site, the fraction of cells expressing the associated gene (at least one read count) was calculated.

#### Total RNA-seq from bulk mouse tissues

RNA extraction was performed on fresh-frozen tissues using the RNAdvance Tissue Total RNA Isolation Kit (A32646, Beckman Coulter) according to manufacturer’s instructions. In short, the tissues were homogenised in Lysis LBE + Proteinase K using the TissueLyzer III (Qiagen) for 3 minutes at 25Hz and lysed at 37°C for 30 minutes. The cell pellets were dissolved in Lysis LBE + Proteinase K and lysed at 37°C for 30 minutes. Automated extraction including DNase treatment was performed on a Biomek i7 (Beckman Coulter). An additional DNase treatment with DNase I (AM2224, Ambion, ThermoFisher Scientific) followed by 2.2X beads:sample ratio bead wash with RNA-Clean XP beads (A63987, Beckman Coulter) was performed post extraction to remove remaining genomic DNA. RNA was quantified on a Fragment Analyzer (Agilent) using the RNA Standard Sensitivity kit (DNF-471, Agilent), and RQN values were obtained.

Library preparation was performed on a Tecan Fluent (Tecan) using the KAPA RNA HyperPrep Kit with RiboErase (HMR) (8098140702, Roche); 400ng of total RNA were used as input. Obtained libraries were quantified on a Fragment Analyzer (Agilent) using the NGS Fragment Standard Sensitivity kit (DNF-473, Agilent) and pooled equimolarly using the Tecan Fluent (Tecan). The final pool was quantified on a Qubit (ThermoFisher Scientific) using the Qubit 1X dsDNA High Sensitivity (HS) kit (Q33231, Invitrogen, ThermoFisher Scientific), before diluting and sequencing on a NextSeq2000 (Illumina). Paired-end sequencing with 51bp was performed using a P4 XLEAP 100 cycle kit (20100994, Illumina).

Fastq files were collected and read quality was assessed using FastQC (v0.12.1)^100^, RNAseQC (2.3.5)^101^ and samtools stats (v1.18)^102^. Quality control (QC) metrics for RNAseQC were based on a STAR (v2.7.11a)^103^ alignment against the mouse genome (GRCm39, Gencode vM32). Next, QC metrics were summarized using MultiQC (v1.17)^104^. Sequencing adapters were then trimmed from the remaining libraries using FastP (v0.23.4)^105^. A mouse transcriptome index consisting of Gencode entries was generated and reads were mapped to the index using Salmon (v1.10.1)^106^. The bioinformatics workflow was organized using Nextflow workflow management system (v23.10)^107^ and the Bioconda software repository^108^.

#### DNA methylation analysis of published data sets

Methylation signal in bedgraph format was downloaded from NCBI GEO; four mESC samples from two different studies (GSM6369000, GSM6369001, GSM5138751 and GSM5138752)^66, 67^ and ENCODE (https://www.encodeproject.org) WGBS data from mouse spleen, lungs, liver, colon, kidney and heart (ENCFF318OTG, ENCFF322GLW, ENCFF646MMY, ENCFF942CDV, ENCFF590HWX, ENCFF745UHD, ENCFF331NGZ, ENCFF350JPL, ENCFF368EPE and ENCFF443JGV)^68–72^. Average methylation was calculated over 1kb windows centered on the regions of interest with BEDTools^109^.

#### Animal studies

All animal experiments were approved by *the AstraZeneca internal committee for animal studies* as well as *the Gothenburg Ethics Committee for Experimental Animals* (license number 2194-2019), compliant with EU directives on the protection of animals used for scientific purposes. C57BL/6N mice (Charles River) were group-housed in a temperature (21 ± 2 °C) and humidity (55 ± 15%) controlled room with a 12:12-h light:dark cycle. Food and tap water were provided ad libitum. For cage bedding and enrichments, aspen chips, shredded paper, gnaw sticks and a paper house were used.

For generation of the inducible *Pcsk9*-editing mouse models, two different DNA targeting vectors were generated for homologous targeting of the Rosa26 locus^110^ in mouse embryonic stem cells (mESC). The vectors contain two different expression units – one for the U6-driven constitutive expression of either the promiscuous gP or the specific gMH guide RNA and one Tet-on 3G-driven cassette for the doxycycline-inducible expression of SpCas9 (see Extended Data Fig. 6). To minimize dysregulation, each expression unit is flanked by double 1.2 kb insulators derived from the chicken beta globin locus^111^. Homology sequences for targeting the mouse Rosa26 locus surround the expression cassettes. For positive selection, a floxed, self-excising Neomycin resistance cassette is included in the vector. To induce *in vivo* deletion of the cassette, Cre recombinase is expressed from a 698 bp testis-specific angiotensin converting enzyme (tAce) promoter, which is active during early spermatogenesis^112^.

The DNA constructs were electroporated into Primogenix C57BL6/N (PrX) ES cells. Neo-resistant clones were screened by PCR for correct targeting. Two clones per cell line were sent to Cergentis for validation by Targeted Locus Amplification. One validated clone per line was injected into Balb/cAnCrl blastocysts to generate chimeric mice. Chimeric males were bred to C57BL/6N Crl females and black-coated offspring were genotyped for carrying the modified inducible R26 U6 gP CAG Tet-on SpCas9 or R26 U6 gMH CAG Tet-on SpCas9 allele.

Three different genotypes were used in the gene editing study: (1) R26-U6-gP-Tet-on-SpCas9 mice expressing the promiscuous gP alongside inducible SpCas9; (2) R26-U6-gMH-Tet-on-SpCas9 mice expressing the specific gMH alongside inducible SpCas9; and (3) wild-type littermates. Both male and female mice were used in this study. Animals were randomly assigned to either the control or treatment cohorts with n=4 mice for each experimental group/genotype. For induction of *Pcsk9 in vivo* editing, treatment groups received an initial doxycycline dose of 50 mg/kg bodyweight via *per os* (PO) administration, followed by long-term administration of doxycycline at 0.2 mg/ml in the drinking water for 12 days. The wellbeing and body weight of all mice was monitored on a daily basis and any decrease in weight was counteracted by addition of high-nutrient wet food supply. During termination, the different organs (brain, colon, heart, kidney, liver, lungs, pancreas, spleen) were taken out and snap frozen in liquid nitrogen prior to storage at -80 °C. DNA and RNA for downstream analysis were isolated from the snap-frozen tissues using the AllPrep 96 DNA/RNA kit (QIAGEN) according to manufacturer’s instructions. Editing analysis was performed as described above.

### Differentiation of mESC-derived cell types

#### Astrocyte differentiation

Generation of mouse embryonic stem cell (mESC)-derived astrocytes was done as previously published^113^. Briefly, R26-U6-gP-Tet-on-SpCas9 mESCs were trypsinized at 60-80% confluency, counted and seeded (Day 0) at 500,000 cells/mL onto low attachment petri dishes in Advanced DMEM-F12:Neurobasal media (ADFNK) medium composed of advanced DMEM/F12–neurobasal (1:1; Gibco), 10% knockout serum replacement (KSR; Gibco), 1% L-glutamine (Gibco), 100 μM 2-mercaptoethanol (Gibco) and 1% Pen/Strep (Gibco). Cells were maintained at 37 °C in 95% O2, 5% CO2, for the formation of embryoid bodies (EBs). The medium was exchanged and supplemented with 1 μM retinoic acid (RA) (Sigma Aldrich) on Day 2. On Day 5, the medium was exchanged for ADFNK without RA. On Day 6, 20 - 50 EBs were seeded onto 0.1% gelatin (StemCell) coated 24 well culture plates containing astrocyte differentiation media composed of advanced DMEM/F12 (Gibco), 2% fetal bovin serum (Gibco), 200 mM L-glutamine (Gibco), 100 µM b-mercaptoethanol (Gibco), 1% Pen/Strep (Gibco) and 50 µg/ml heparin (Sigma). Cells were cultured for 28 days with media change every 2 days. On day 28, cells were treated with 2 µg/ml Doxycycline (ThermoFisher) for 7 days to induce SpCas9 expression.

#### Hepatocyte differentiation

Hepatic differentiation was conducted following a protocol adapted from Pauwelyn et al.^114^. R26-U6-gP-Tet-on-SpCas9 mESC were trypsinized, and 40,000 cells were seeded on 2% matrigel (BD) coated 24-well plates in basal differentiation medium (BDM; 60% DMEM-low glucose (Gibco), 40% DMEM/F12 (Gibco), 0.25x linoleic acid-albumin from bovine serum albumin (Sigma), 0.25x insulin-transferrin-selenium (Sigma), 1% Pen/Strep (Gibco), 0.1 µM L-ascorbic acid (Sigma), 0.001 µM dexamethasone (Sigma), and 110 µM 2-mercaptoethanol (Gibco)). Cytokine cocktails (R&D Systems) were added to BDM as follows: day 0 - 2, 2% FBS (Gibco) was added and supplemented with 100 ng/ml activin-A and 100 ng/ml Wnt3a; day 3 - 6, FBS was reduced to 0.5% and supplemented with 100 ng/ml activin-A and 100 ng/ml Wnt3a; day 6 - 10, 0.5% FBS was added and supplemented with 10 ng/ml FGF2 and 50 ng/ml BMP4; from day 10 - 14, 0.5% FBS was added and supplemented with 25 ng/ml FGF8b, 50 ng/ml FGF1 and 10 ng/ml FGF4 and from day 14 onwards, 0.5% FBS with 20 ng/ml HGF and 100 ng/ml follistatin. On day 26, cells were treated with doxycycline (ThermoFisher) for 7 days to induce SpCas9 expression and harvested afterwards for DNA and RNA extraction. Media was exchanged every two days.

#### Cardiomyocyte differentiation

Differentiation of R26-U6-gP-Tet-on-SpCas9 mESCs into cardiomyocytes was performed as described previously^115^ with some modifications. mESC were seeded onto 24-well gelatin-coated plates at a cell density of 50,000 cells/well for 8h in IMDEM (Gibco) medium plus LIF (Merk) before transitioning to Day 0 medium. Day 0 medium consisted of IMDM/Ham’s F12 (Invitrogen), N2 supplement (Gibco), B27 supplement (Gibco), 10% bovine serum albumin (BSA) (Sigma), L-glutamine (Gibco), 1% Pen/Strep (Gibco), 0.5 mM ascorbic acid (Sigma), and 450 µM monothioglycerol (MTG, Sigma) for 1 day. For mesodermal induction, 8 ng/ml activin A (R&D Systems), 5 ng/ml VEGF (R&D Systems) and 0.5 ng/ml BMP4 (R&D Systems) were used for 2 days. Cardiac specification was induced with StemPro-34 SF medium (Gibco) supplemented with 2 mM L-glutamine (Gibco), 0.5 mM ascorbic acid (Sigma), 5 ng/ml VEGF (R&D Systems), 10 ng/ml bFGF (R&D Systems), and 50 ng/ml FGF10 (R&D Systems) until the end of differentiation. Twelve days after differentiation, cardiomyocytes were treated with 2 µg/ml doxycycline (ThermoFisher) for 5 days to induce SpCas9 expression and then collected for DNA and RNA extraction. Medium was changed every other day, and cells were washed twice with phosphate buffered saline (PBS, Gibco) when changing to different media.

#### Cell type-marker analysis

To assess differentiation success, expression of cell type-specific markers was analyzed by real-time RT qPCR. To this end, total RNA was extracted from harvested cells using the AllPrep DNA/RNA Mini Kit (QIAGEN) and subsequently converted to cDNA with the High-Capacity cDNA Reverse Transcription Kit (ThermoFisher), according to the manufacturers’ instructions. SYBR-Green gene expression assay kits (Applied Biosystems) were used for the quantification of cell type markers (Supplementary Table 8). Glyceraldehydes-3-phosphate dehydrogenase (*Gapdh*) was used as endogenous control gene for normalization. All qPCR experiments were run on a QuantStudio 7 Flex system (ThermoFisher) using the following protocol: 50 °C for 2 min, 95 °C for 10 min, 40 cycles of 95 °C for 15 sec and 60 °C for 1 min.

### SpCas9 expression analysis

For SpCas9 expression analysis, RNA isolated from mouse tissues was reverse-transcribed into cDNA using the High-Capacity cDNA Reverse Transcription Kit (ThermoFisher), according to the manufacturer’s instructions. qPCR analysis was done using TATAA SYBR GrandMaster Mix ROX (TATAA Biocenter AB) and custom designed primers (see Supplementary Table 8) on a QuantStudio 7 Flex system (ThermoFisher) using the following protocol: 95 °C for 30 sec, 40 cycles of 95 °C for 5 sec, 60 °C for 30 sec and 72 °C for 20 sec, followed by melting curve analysis. Absolute abundance of SpCas9 RNA was calculated from an input standard curve generated from synthetic RNA-derived cDNA.

### Data analysis and presentation

Data visualization and statistical analysis were done with GraphPad Prism 10 (GraphPad Software, Inc.). Detailed information on statistical tests, sample sizes, and P values are found in the figure legends. P ≤ 0.05 was considered statistically significant. Illustrations were created with BioRender.com.

## Acknowledgements

We would like to thank the AstraZeneca NGS & Transcriptomics team for supporting the targeted amplicon sequencing experiments. We are grateful to Ben Keith, Ramy Elgendy, Petr Volkov, Graham Belfield, Sasa Svikovic and Yousra Yahia for the discussions on this work and to Liselotte Andersson for her support on animal work. This project was funded by the AstraZeneca Postdoc program. A.M. and M.M.L.G. are former AstraZeneca postdoc program fellows. M.S. is part of the AstraZeneca Graduate Program.

## Author contributions

A.M., M.M. and P.Ak. conceptualized the study. A.M., N.S., M.M.L.G, M.S., and P.An. performed *in vitro* cell experiments, sequencing library preparations, editing analyses. A.L.L., K.M., J.F. and K.M.B. performed embryo electroporations, mESC line and mouse line generations. M.S. performed WGA experiments. J.C. performed scATAC-seq and scRNA-seq experiments. J.L. and J.L.T. performed sequencing experiments. D.J. performed long-read sequencing. I.D. performed bulk RNA-seq. M.F., L.W., B.S. and K.N. performed bioinformatic analyses. G.K. performed PASTA translocation analyses. A.M., A.E., J.H. and M.P. performed *in vivo* editing experiments. A.J., M.B. and M.M. facilitated collaborative work and provided scientific guidance. A.M., N.S., and P.Ak. prepared the manuscript with input from all authors. P.Ak. supervised the study.

### Data availability

DNA sequencing data are available via the NCBI Sequence Read Archive (PRJNA1461838).

## Extended Data

**Extended Data Fig. 1.**
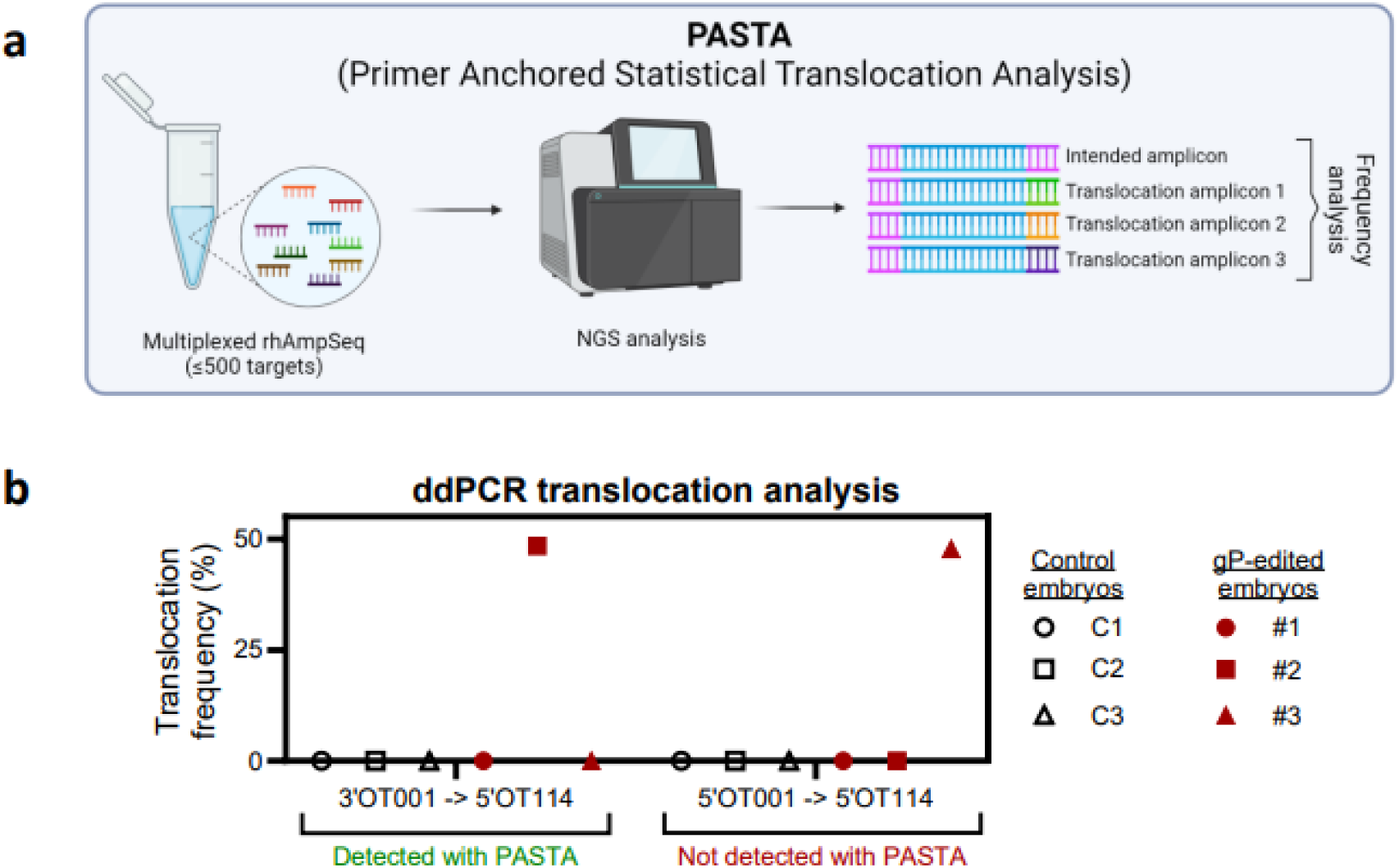
Translocation analysis with PASTA and ddPCR. a. Schematic of PASTA translocation analysis. The multiplexed nature of rhAmpSeq gives rise to translocation product amplicons during library preparation in addition to the expected amplicons of off-target sites, which allows frequency analysis of the detected translocations using the rhAmpSeq sequencing data. Created in BioRender. Madsen, A. (2026) https://BioRender.com/r8ngkyt. b. Targeted translocation analysis in mouse embryos by ddPCR. Detectability of the indicated translocations by PASTA is indicated at the bottom of the graph.

**Extended Data Fig. 2.**
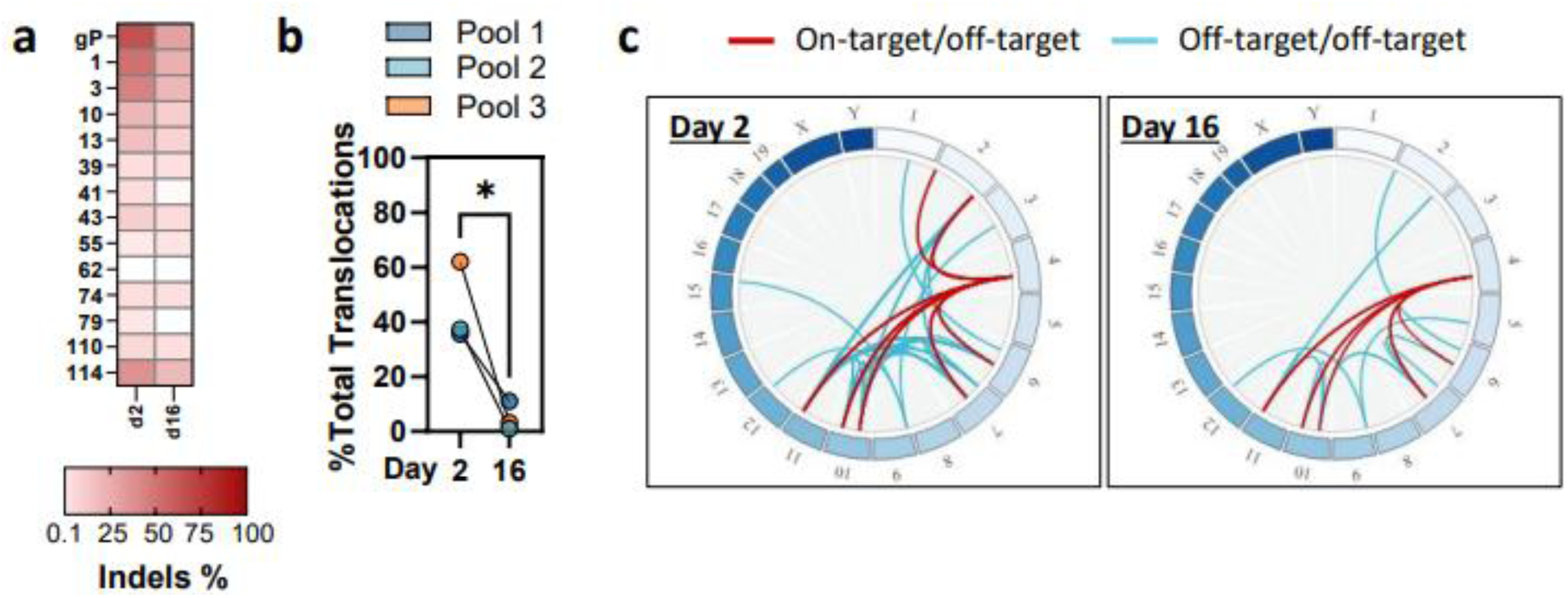
Changes in off-target editing and translocations in mESC pools over time. a. Heatmap showing the percentage of indels detected at the indicated promiscuous gP sgRNA on-/off-target sites in pools of bulk-edited mESCs. The pools show mean editing of n=3 replicate pools 2 days and 16 days after editing. All sites with >0.1% editing in at least one of the pools are shown. b. Dot plots showing total translocation burden in the gP-edited mESC pools over time, calculated as the sum of the frequencies of all detected translocations for each of the three individual cell pools on day 2 and day 16 after electroporation. % Total translocations is calculated as the sum of the frequencies of all detected translocations at a given time point. Paired t-test, * = p<0.05. c. Circos plots showing translocations detected by PASTA in the mESC pools on day 2 and day 16, respectively. Red lines indicate translocations between the on-target and off-target sites, blue lines indicate translocations between off-target sites.

**Extended Data Fig. 3.**
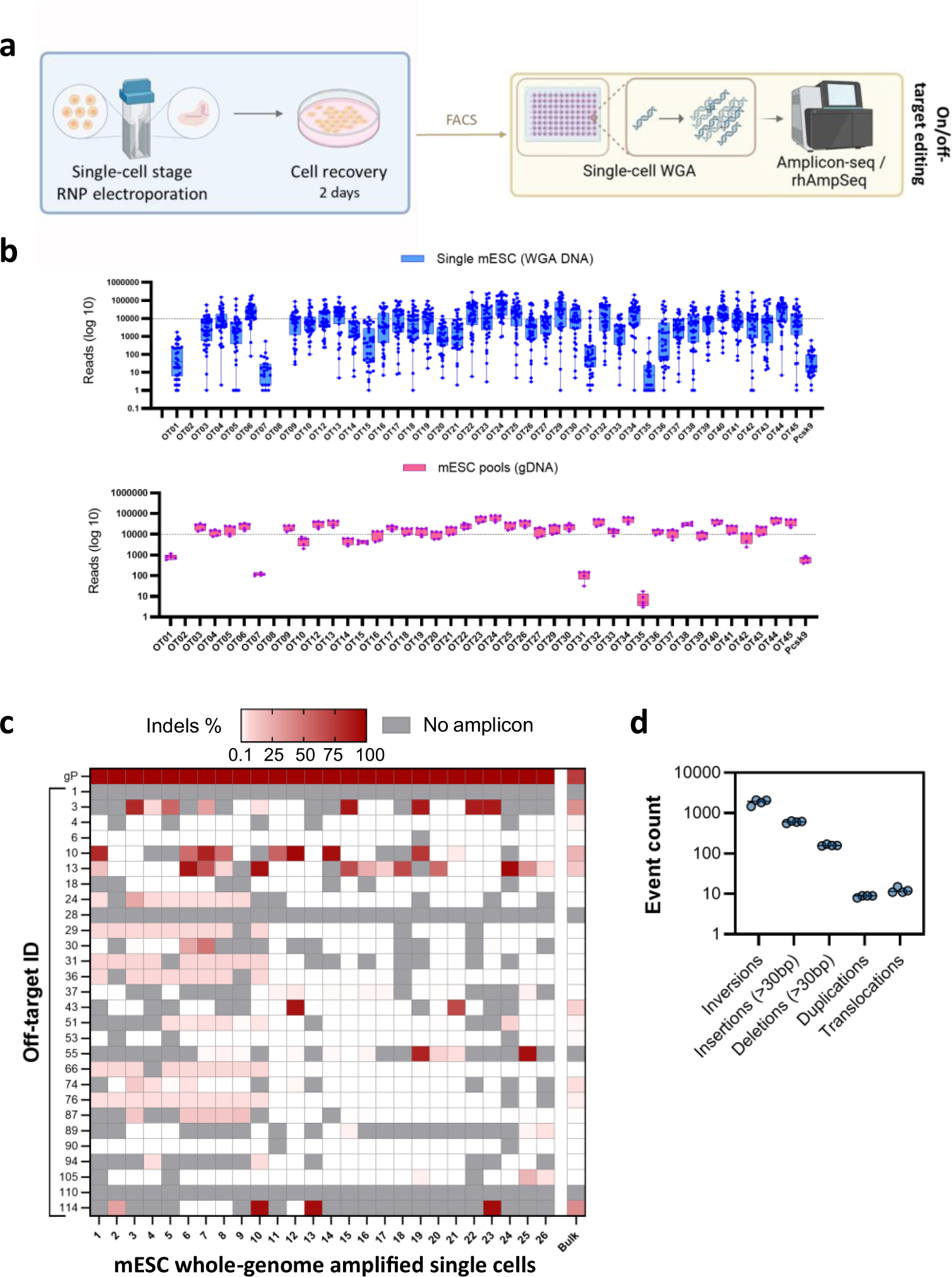
**Whole-genome amplification of edited single cells.** a. Overview of the experimental workflow for single-cell editing analyses in mESCs using single-cell whole-genome amplification instead of clonal expansion. Created in BioRender. Madsen, A. (2026) https://BioRender.com/2zywinr. b. Inter-sample variation of site coverage after rhAmpSeq using WGA DNA as input (upper panel) vs. using genomic DNA (bottom panel). The dotted line indicates a coverage of 10,000x, which is needed to reliably detect mutations with a sensitivity of 0.1% of editing. c. Heatmap showing the percentage of indels detected at the indicated promiscuous gP sgRNA on-/off-target sites in whole-genome amplified single-cell edited mESCs and a pool of bulk-edited mESCs. d. Large structural variants detected by whole-genome long-read sequencing of whole-genome amplified mESCs. Each data point represents a biological replicate.

**Extended Data Fig. 4.**
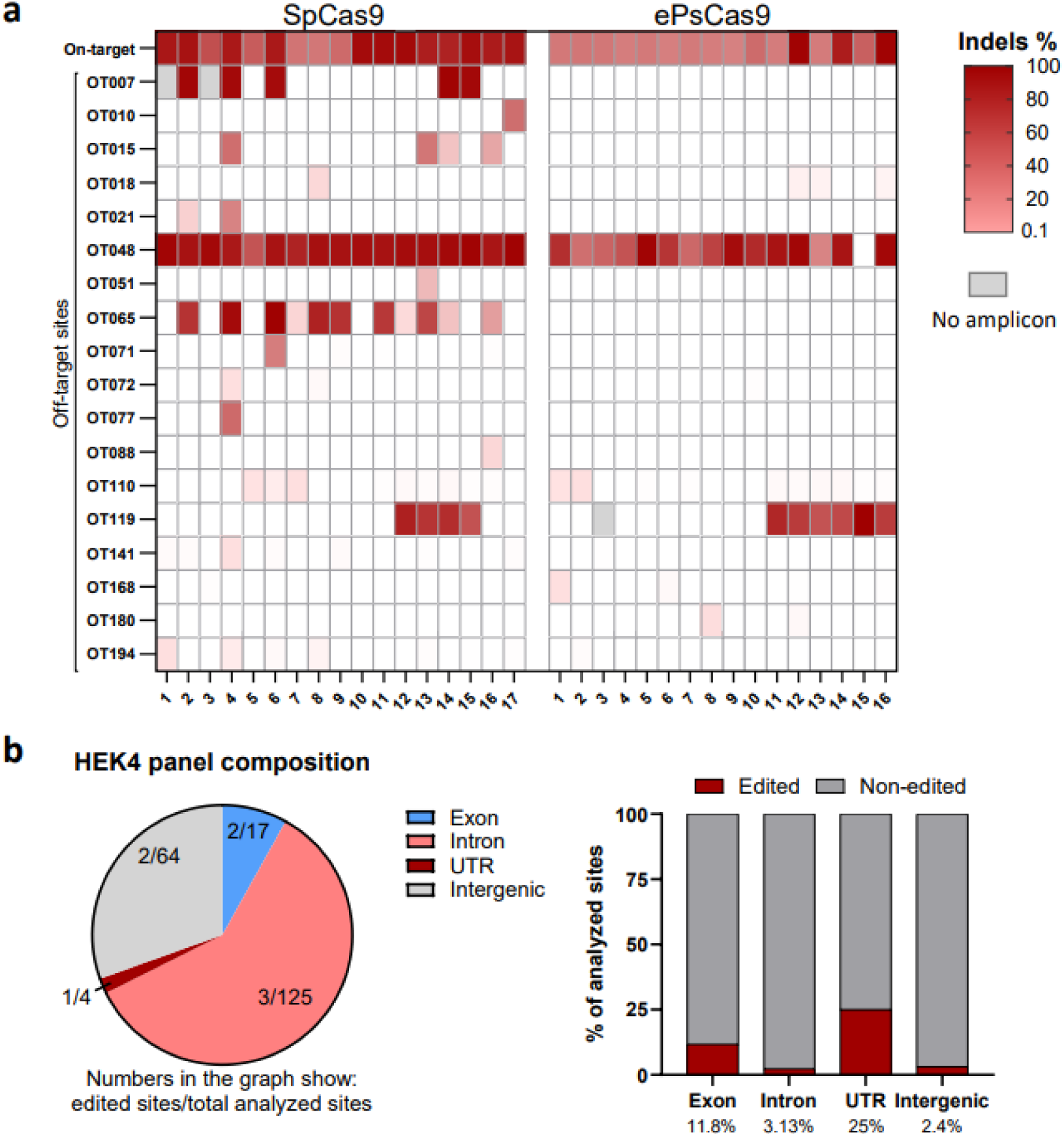
Off-target analysis in single cell-edited HSPCs. a. Heatmap showing off-target editing in expanded single cell-edited HSPCs after editing with SpCas9 or ePsCas9 and the HEK4 sgRNA. Each column represents a single-cell clone. b. Overview of the HEK4 rhAmpSeq panel composition regarding coverage of exonic, intronic, UTR and intergenic regions, as well as the fraction of edited sites for each sub-group.

**Extended Data Fig. 5.**
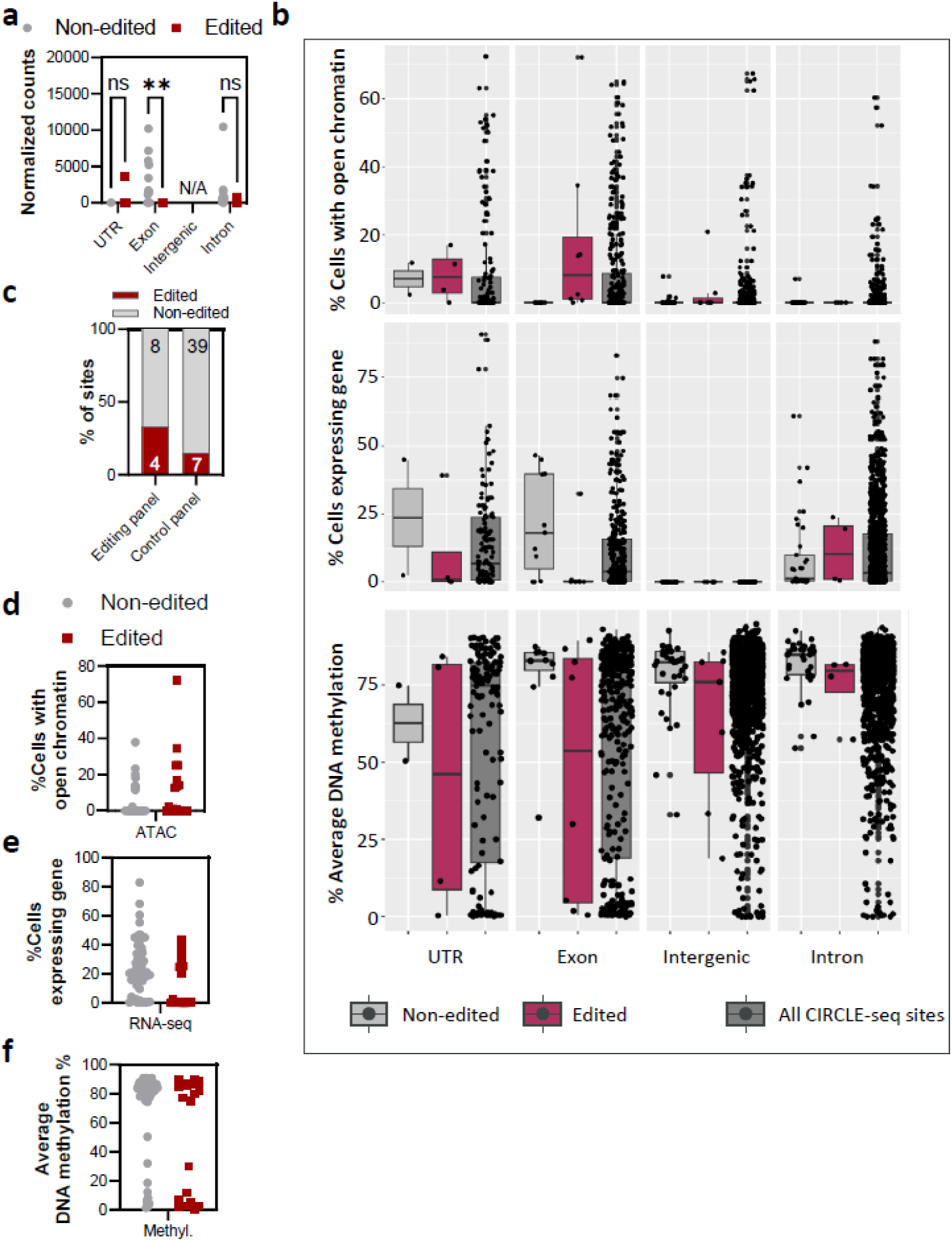
Validation of observations from scATAC-seq/scRNA-seq experiments. a. Bulk RNA-sequencing of mESCs to validate observed correlations between editing and gene expression for exonic sites from scRNA-seq results. b. Comparison of distribution of open chromatin (upper panel), gene expression (middle panel) and DNA methylation (lower panel) in the analyzed sites vs. all CIRCLE-seq-identified sites. c. Overview of gP validation rhAmpSeq panels. The “Editing panel” included sites that were considered likely to be edited according to the proposed prediction features, whereas the “Control panel” included sites unlikely to be edited. The graph shows the number of sites for both panels that showed editing or remained unedited. d. Analysis of scATAC-seq data from mock-electroporated mESCs based on functional region of the respective gP off-target site from the replication panels. Plot showing percentage of cells with open chromatin at sites showing editing vs. sites that remain non-edited. e. Analysis of scRNA-seq data from mock-electroporated mESCs based on functional region of the respective gP off-target site from the replication panels. Plot showing percentage of cells expressing the off-target-associated gene at sites showing editing vs. sites that remain non-edited. f. DNA methylation analysis from whole-genome bisulfite sequencing data^66, 67^ at gP off-target sites from the replication panels showing editing vs. sites that remain non-edited. Plot showing the average methylation percentage of a region ±500 bp around each sgRNA off-target site.

**Extended Data Fig. 6.**
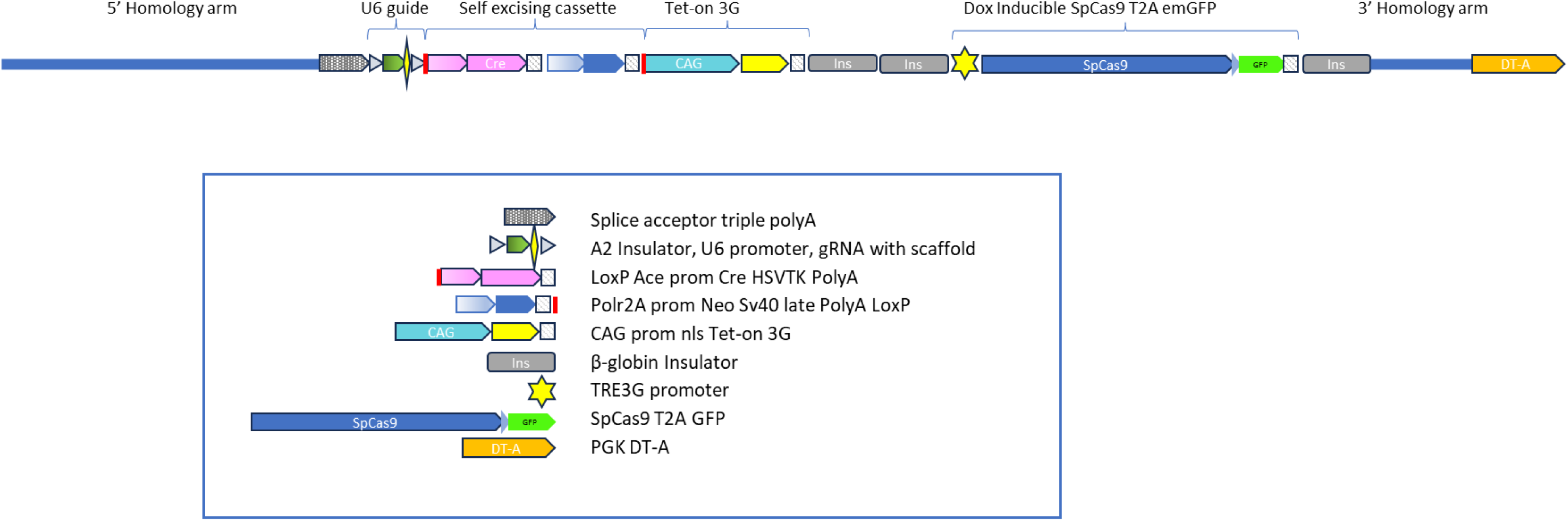
Schematic of the targeting vector “R26-U6-gP/gMH-Tet-on-SpCas9” that was used for generation of editing-inducible mESC lines.

**Extended Data Fig. 7.**
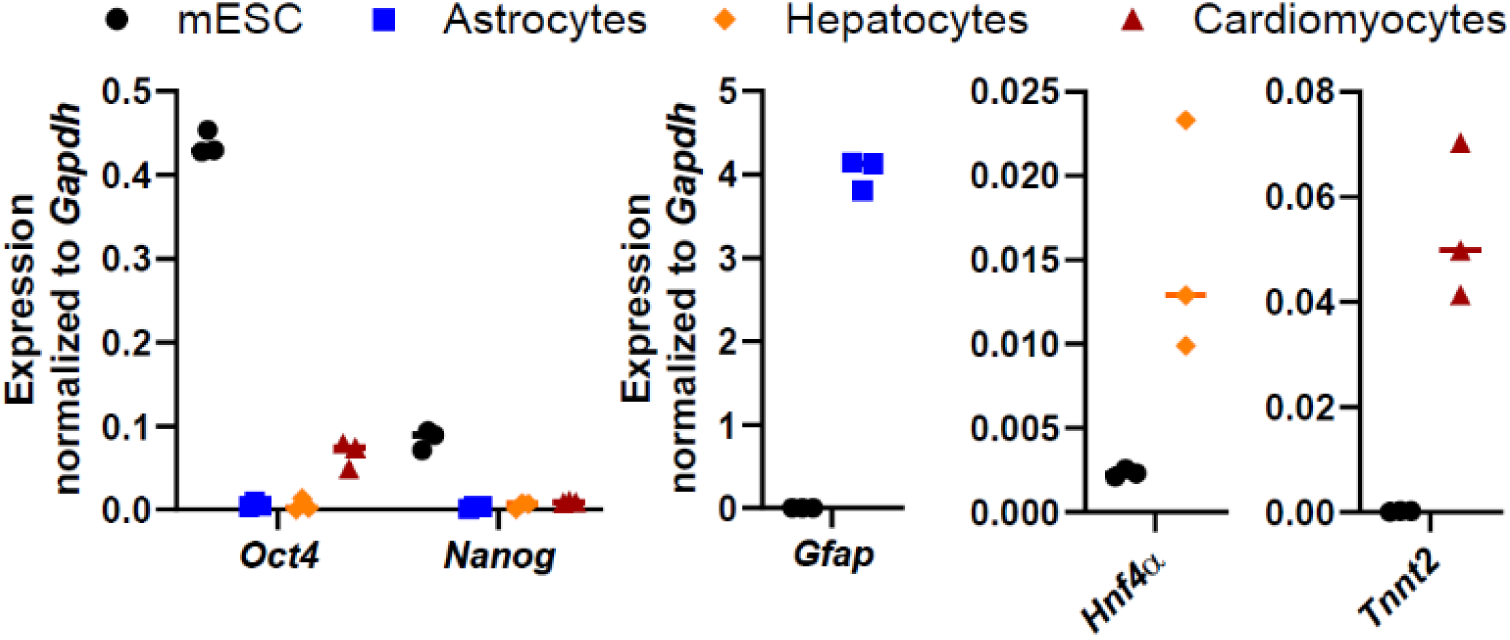
Validation of successful mESC differentiation by cell marker expression analysis. Differentiated mESCs showed a reduced expression of the pluripotency markers *Oct4* and *Nanog*, as well as increased expression of the respective cell type-specific markers, i.e. *Gfap* for astrocytes, *Hnf4a* for hepatocytes and *Tnnt2* for cardiomyocytes. Expression of all marker genes was normalized to the endogenous control gene *Gapdh*.

**Extended Data Fig. 8.**
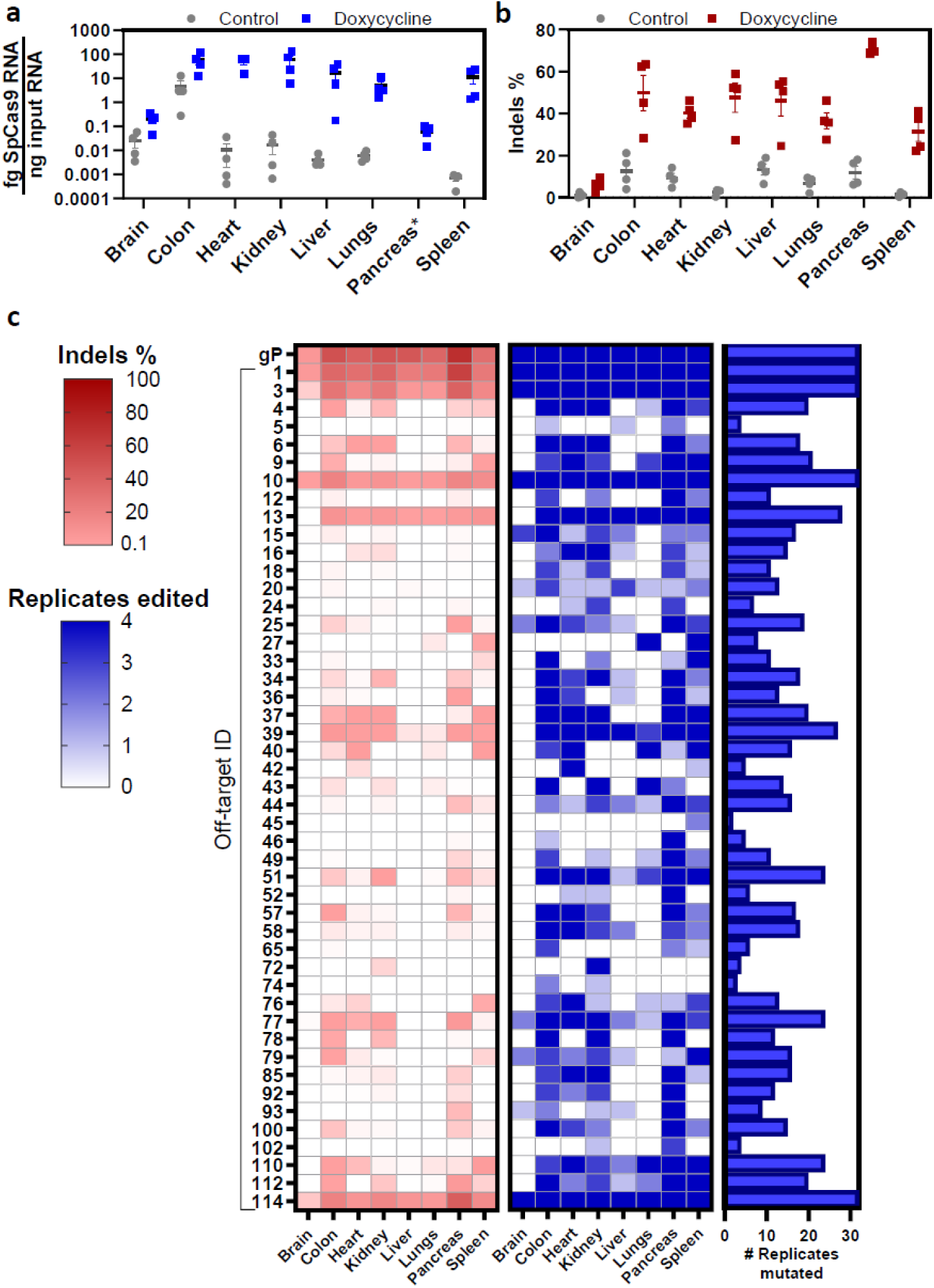
***In vivo* gP editing.** a. SpCas9 RNA expression in the different organs with or without doxycycline treatment. Graph shows mean±SEM SpCas9 RNA (fg) per ng input RNA for n=4 organ replicates. *Note that RNA quality for pancreas samples was very low compared to the other organs due to the high levels of endogenous RNases. b. Percentage of indels detected at the *Pcsk9* on-target site in the different organs with or without doxycycline treatment. Graph shows mean±SEM of editing % for n=4 organ replicates. c. Reproducibility of editing results within organ replicates in vivo. The first heat map shows the mean percentage of indels detected in n=4 organ replicates for sites that exhibit >0.1 % indels in at least one organ. The second heat map shows number of replicates (0-4) that exhibit >0.1% indels at the indicated sites within each organ group. The Waterfall plot on the right shows the total number of replicates edited across all groups.

**Extended Data Fig. 9.**
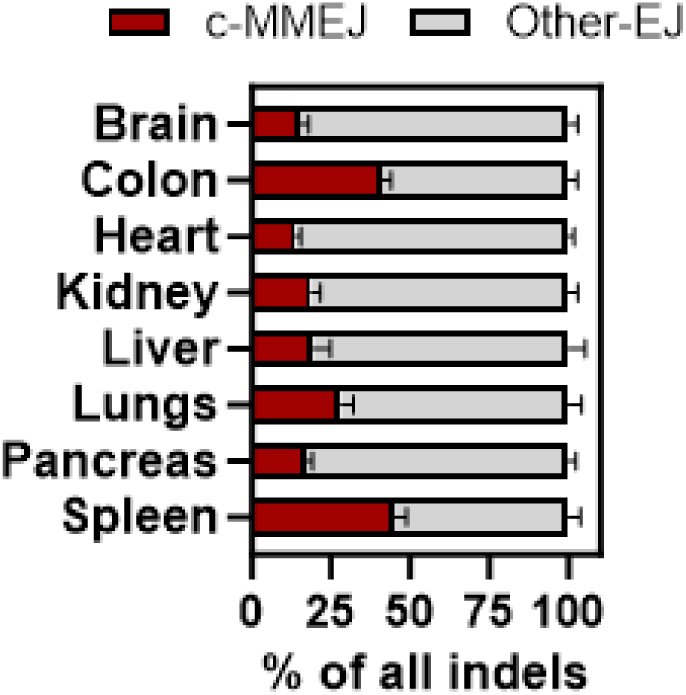
Comparison of DNA repair mechanisms in the different organs at all sites that exhibit >0.1% indels in the respective organs upon doxycycline-induced SpCas9 expression with gP. n=4 organ replicates. c-MMEJ = micro-homology mediated end-joining; Other-EJ = other end-joining.

**Extended Data Fig. 10.**
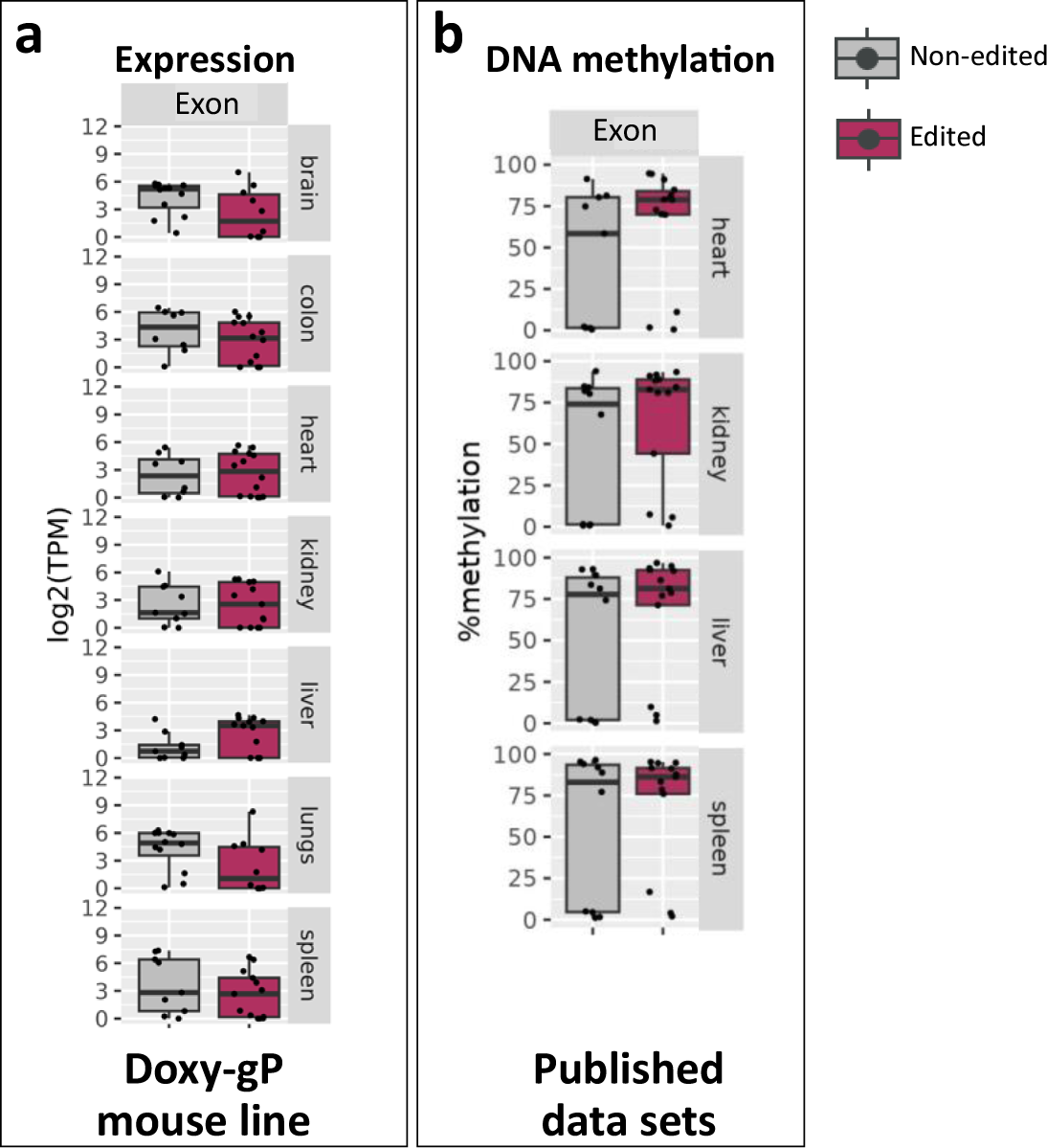
**Correlation of gene expression and DNA methylation in mouse tissues with the observed gP off-target editing after SpCas9 induction.** a. RNA-sequencing analysis of gene expression in the doxy-gP mouse line. b. Analysis of publicly available DNA methylation data sets from an unrelated mouse line^68–72^.

**Extended Data Fig. 11.**
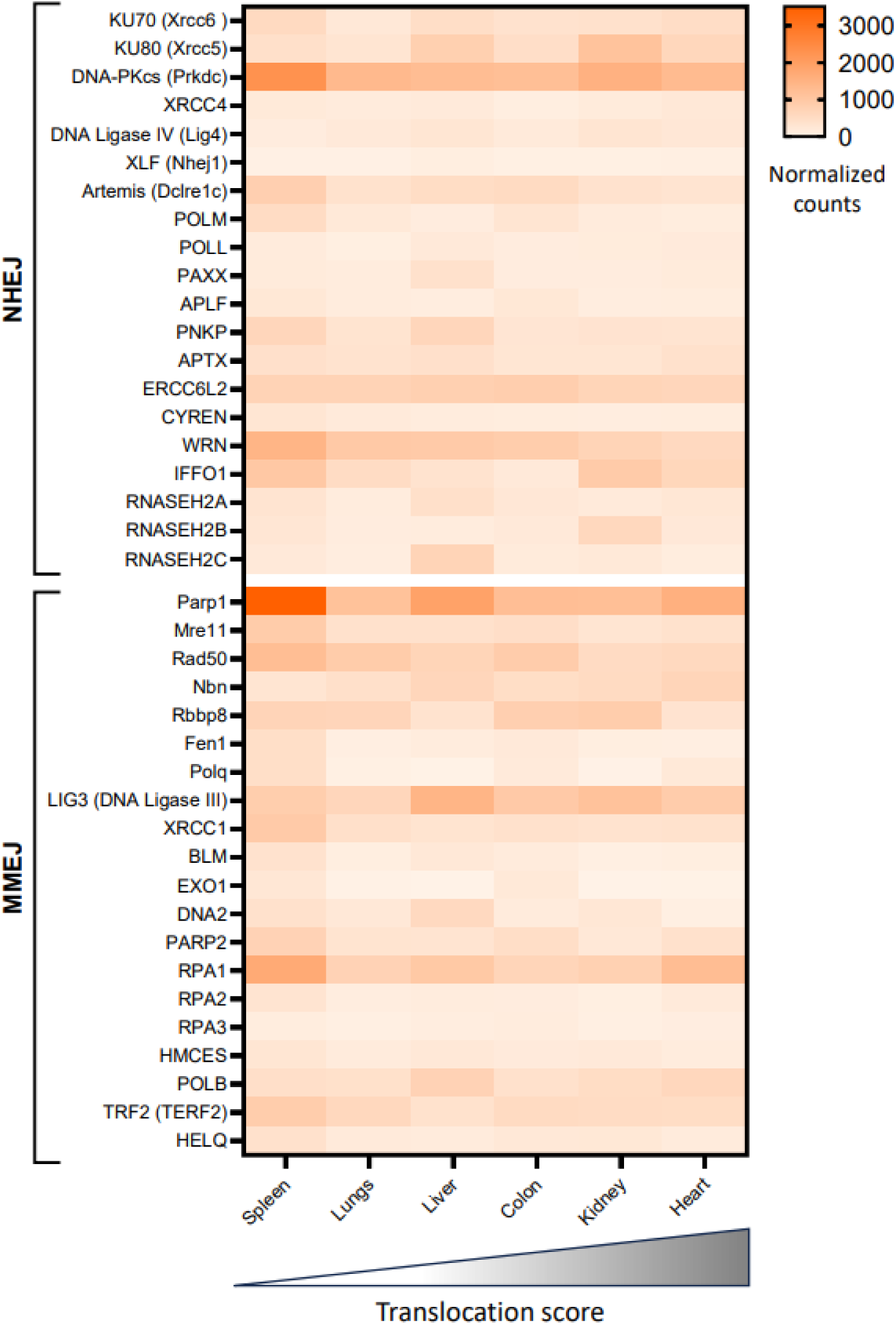
Heatmap showing an overview of DNA repair gene expression in the different mouse organs. The color scale shows the number of normalized read counts from RNA-sequencing. The mean of n=4 animals is shown for each organ. Organs on the x-axis are sorted according to their translocation score from lowest to highest.

**Extended Data Fig. 12.**
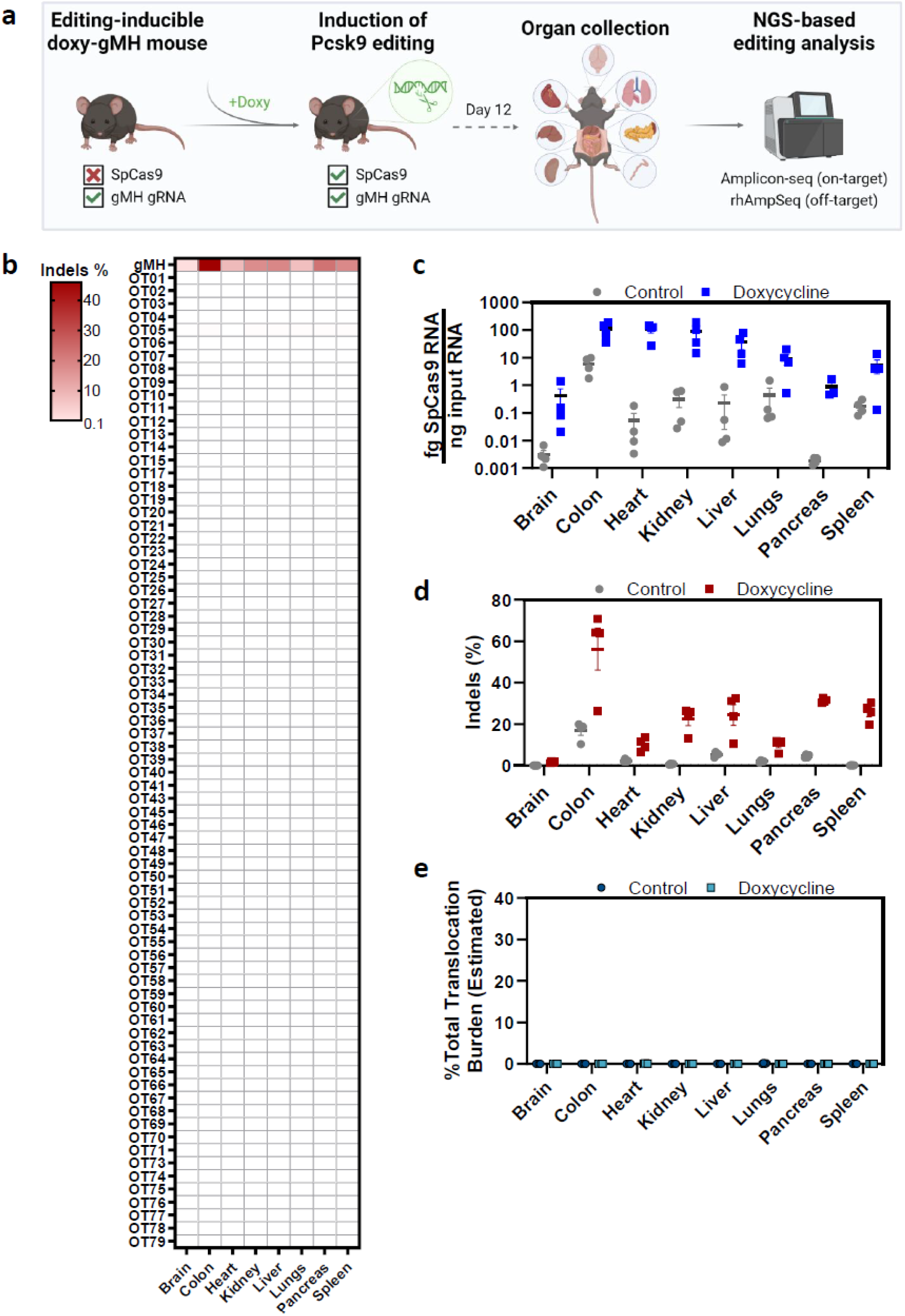
**Analysis of gMH-mediated editing outcome in different organs *in vivo*.** a. Overview of the *in vivo* editing experiments using a doxycycline-inducible SpCas9-gMH (doxy-gMH) mouse model. Created in BioRender. Madsen, A. (2026) https://BioRender.com/71p3urb. b. Heat map showing the mean percentage of indels detected in different organs upon editing induction by doxycycline treatment. n=4 organ replicates. c. SpCas9 RNA expression in the different organs with or without doxycycline treatment. Graph shows mean±SEM SpCas9 RNA (fg) per ng input RNA for n=4 organ replicates. d. Percentage of indels detected at the *Pcsk9* on-target site in the different organs with or without doxycycline treatment. Graph shows mean±SEM of indels % for n=4 organ replicates. e. Comparison of translocation burden in the different organs. Shown is the total translocation burden (i.e. sum of frequency of all detected translocations per organ). No significant occurrence of translocations was detected in the different organs after gMH editing.

## Supplementary Tables

**Supplementary Table 1** – rhAmpSeq panel information

**Supplementary Table 2** – rhAmpSeq results single-cell editing, normalized to sequencing controls

**Supplementary Table 3** – Results from PASTA translocation analysis

**Supplementary Table 4** – Results of scATAC-seq/scRNA-seq analysis

**Supplementary Table 5** – Results of DNA methylation analysis

**Supplementary Table 6** – rhAmpSeq results cell types and tissues, normalized to sequencing controls, including statistical analysis of editing

**Supplementary Table 7** – Bulk RNA-seq from mouse tissues and mESCs, normalized counts

**Supplementary Table 8** – Primer and guide information

